# mTOR is a Major Determinant of Chemosensitivity

**DOI:** 10.1101/2022.01.19.476916

**Authors:** Yuanhui Liu, Nancy G. Azizian, Delaney K. Sullivan, Yulin Li

## Abstract

Chemotherapy can eradicate a majority of cancer cells. However, a small population of tumor cells often survives drug treatments through genetic and/or non-genetic mechanisms, leading to tumor recurrence. Here we report a reversible chemoresistance phenotype regulated by the mTOR pathway. Through a genome-wide CRISPR knockout library screen in pancreatic cancer cells treated with chemotherapeutic agents, we have identified mTOR pathway as a prominent determinant of chemosensitivity. Pharmacological suppression of mTOR activity in cancer cells from diverse tissue origins leads to the persistence of a reversibly resistant population, which is otherwise eliminated by chemotherapeutic agents. Conversely, activation of mTOR pathway increases chemosensitivity *in vitro* and *in vivo* and predicts better survival among various human cancers. Persister cells display a senescence phenotype. Inhibition of mTOR does not induce cellular senescence *per se*, but rather promotes the survival of senescent cells through regulation of autophagy and G2/M cell cycle arrest, as revealed by a small molecule chemical library screen. Thus, mTOR plays a causal yet paradoxical role in regulating chemosensitivity; inhibition of the mTOR pathway, while suppressing tumor expansion, facilitates the development of a reversible drug-tolerant senescence state.

## INTRODUCTION

Genotoxic damage-inducing chemotherapy is the mainstay treatment for most cancers. Effective chemotherapeutic agents often kill a large fraction of tumor cells, while sparing a small surviving population, which can resurface as future tumor recurrence. Acquired therapeutic resistance occurs due to genetic and/or non-genetic processes^1, 2^. In the context of acute stress resulting from genotoxic chemotherapy, the initial adaptation is essential for tumor cell survival and may allow for subsequent development of chemoresistance through additional genetic and/or non-genetic mechanisms.

The limited clinical success of mTOR inhibitors questions the efficacy of this class of agents, and the use of mTOR as a *bona fide* cancer therapeutic target. Furthermore, recent findings suggest that mTOR pathway possesses tumor suppressor activity^3, 4^ and low mTOR activity is associated with resistance to chemo- and immunotherapies^5, 6^. Here, we have performed a genome-wide CRISPR knockout screen and identified mTOR pathway as a major determinant of chemosensitivity. We show that mTOR inhibition causally leads to the emergence of a reversible drug-tolerant subpopulation with a senescence phenotype, which is dependent on autophagy and G2/M cell cycle arrest for survival.

## RESULTS

### CRISPR screen identifies mTOR pathway as a regulator of chemosensitivity

To systematically identify the genetic determinants of chemosensitivity, we performed a genome-wide CRISPR knockout screen in a murine pancreatic cancer cell line using two genotoxic chemotherapeutic agents, gemcitabine and selinexor (**Figure 1A**). Gemcitabine is a deoxycytidine analog that inhibits DNA synthesis^7^, and selinexor is an inhibitor of chromosome region maintenance 1/exportin 1 (CRM1/XPO1), required for chromosome segregation and mitotic progression^8, 9^. As wild type p53 may interfere with Cas9 function and confound interpretation of chemosensitivity^10, 11^, we utilized the 4292 cell line, derived from a mouse pancreatic cancer model induced by Kras^G12D^ and p53^R172H^ mutants^12, 13^. 4292 cells expressing Cas9 (4292-Cas9) were transduced with the Brie sgRNA library containing 78,637 sgRNAs targeting 19,674 genes^14^, selected with puromycin, and subsequently treated with the two chemotherapeutic agents for 12 days. The screens were performed using relatively low concentrations of the drugs (20nM for gemcitabine, and 0.33µM for selinexor), in order to identify in the same screen enriched and depleted genes corresponding to negative and positive regulators of chemosensitivity.

**Figure 1.**
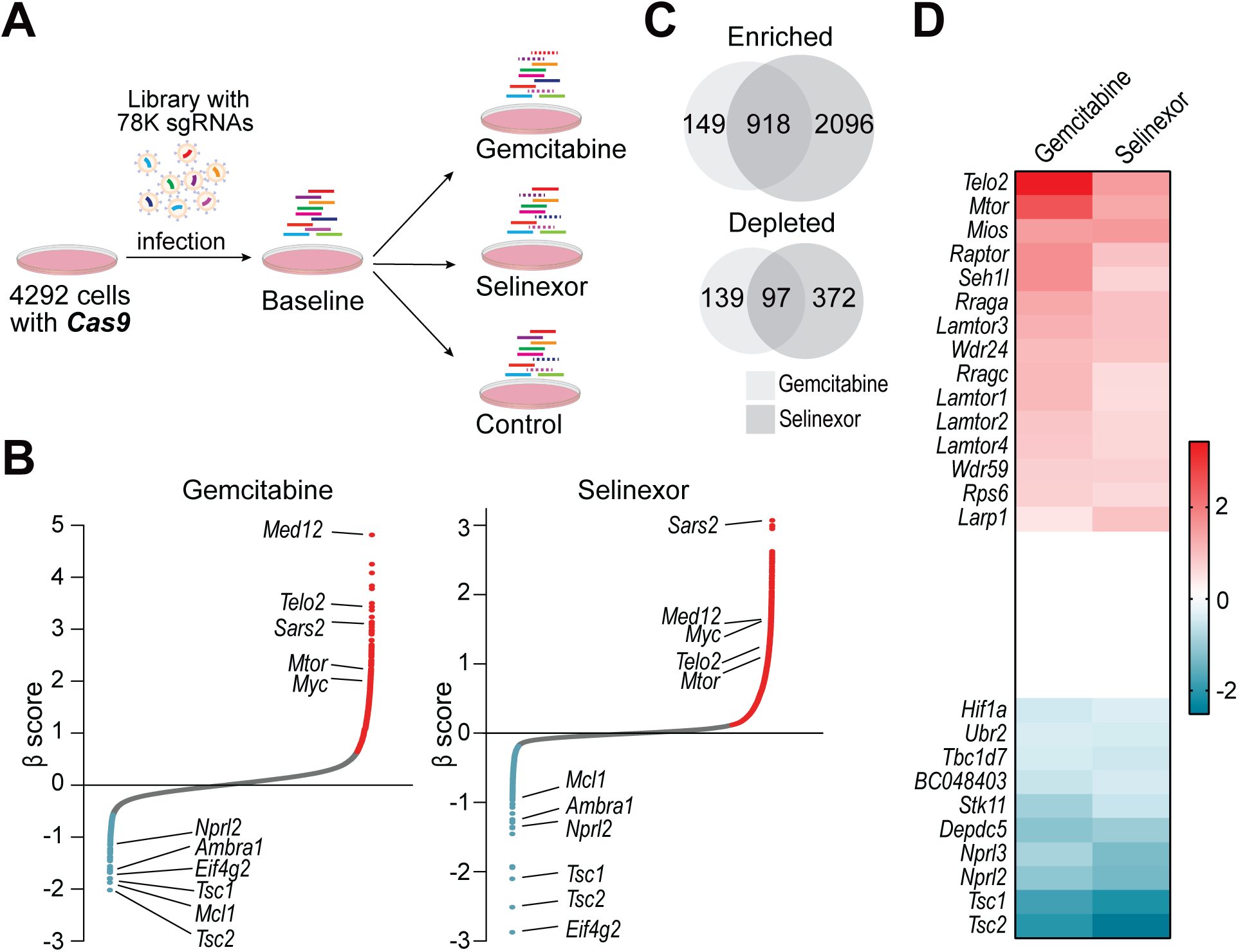
Genome-wide CRISPR screening identifies mTOR pathway as a determinant of chemosensitivity. **A.** CRISPR knockout screens in the 4292 cells. **B.** Gene ranking based on β scores in the two screens. Representative top enriched or depleted genes were labeled on the plots. Red and blue colors indicate significant gene enrichment or depletion (adjusted p < 0.05). **C.** Overlap of significantly enriched or depleted genes in gemcitabine and selinexor screens. **D.** Enrichment of positive regulators, and depletion of negative regulators of mTOR signaling in gemcitabine and selinexor screens.

Following deep sequencing of sgRNA libraries, significantly enriched or depleted genes in each screen were identified based on the β score^15^. Notably, a large fraction of the identified genes was shared by the two screens (**Figures 1B-C; Tables S1A-C**). Gene Ontology (GO) term enrichment analysis was performed using EnrichR^16^ on genes that were significantly changed in both screens. Enriched genes were mainly clustered into protein translation, RNA metabolism and macromolecule biosynthesis, while depleted genes were involved in negative regulation of mTOR signaling and autophagy (**Tables S1D-E**). We specifically examined the β scores of genes involved in mTOR signaling (GO: 0031929), including structural components as well as positive and negative regulators of mTOR complexes. Among the 118 mTOR signaling genes included in the sgRNA library, the top fifteen enriched genes in both screens are exclusively involved in activation of mTOR pathway, including *mTOR*, *Mios*, *Telo2,* and *Raptor;* while top ten depleted genes are negative regulators of mTOR signaling, including *Tsc1*, *Tsc2*, *Stk11,* and *Nprl2/3* (**Figure 1D**). Importantly, most enriched genes are either positive regulators or structural components of the mTORC1, but not the mTORC2 complex. To further validate our findings, we reanalyzed a published CRISPR screen performed in REH leukemia cells carrying a p53 mutation^17^, and confirmed that reduced mTOR pathway activity indeed confers resistance to various DNA damaging agents (**Figure S1**). Hence, our CRISPR screen identifies mTOR pathway as a prominent determinant of chemosensitivity.

### mTOR inhibition promotes the emergence of drug-tolerant persisters

To examine the role of mTOR in regulating chemosensitivity, we compared the efficacy of single agent chemotherapy to chemotherapy plus mTOR inhibitors (mTORi). 4292 cells were treated with single agent gemcitabine or selinexor to determine the concentrations that would completely eradicate tumor cells within seven days. We then treated the cells with each drug as a single agent at the killing concentrations, as well as in combination with rapamycin, and monitored their survival over the course of two weeks. Interestingly, combined treatment, while killing a large fraction of the cells, led to a subpopulation of survivors (**Figure 2A**). Emergence of the drug-tolerant populations was further confirmed using additional rapalogs specific to mTORC1 (temsirolimus and everolimus), as well as mTORC1/2 dual inhibitors (Torin1 and INK128). In contrast, the mTORC2-specific inhibitor, JR-AB2-011, failed to induce the drug-tolerant subpopulation, indicating that the reduced mTORC1, but not mTORC2 activity, leads to chemoresistance (**Figure 2A**). In the presence of drug treatment, surviving tumor cells remained dormant and viable for over three weeks (longest time of observation), herein referred to as “persisters”.

**Figure 2.**
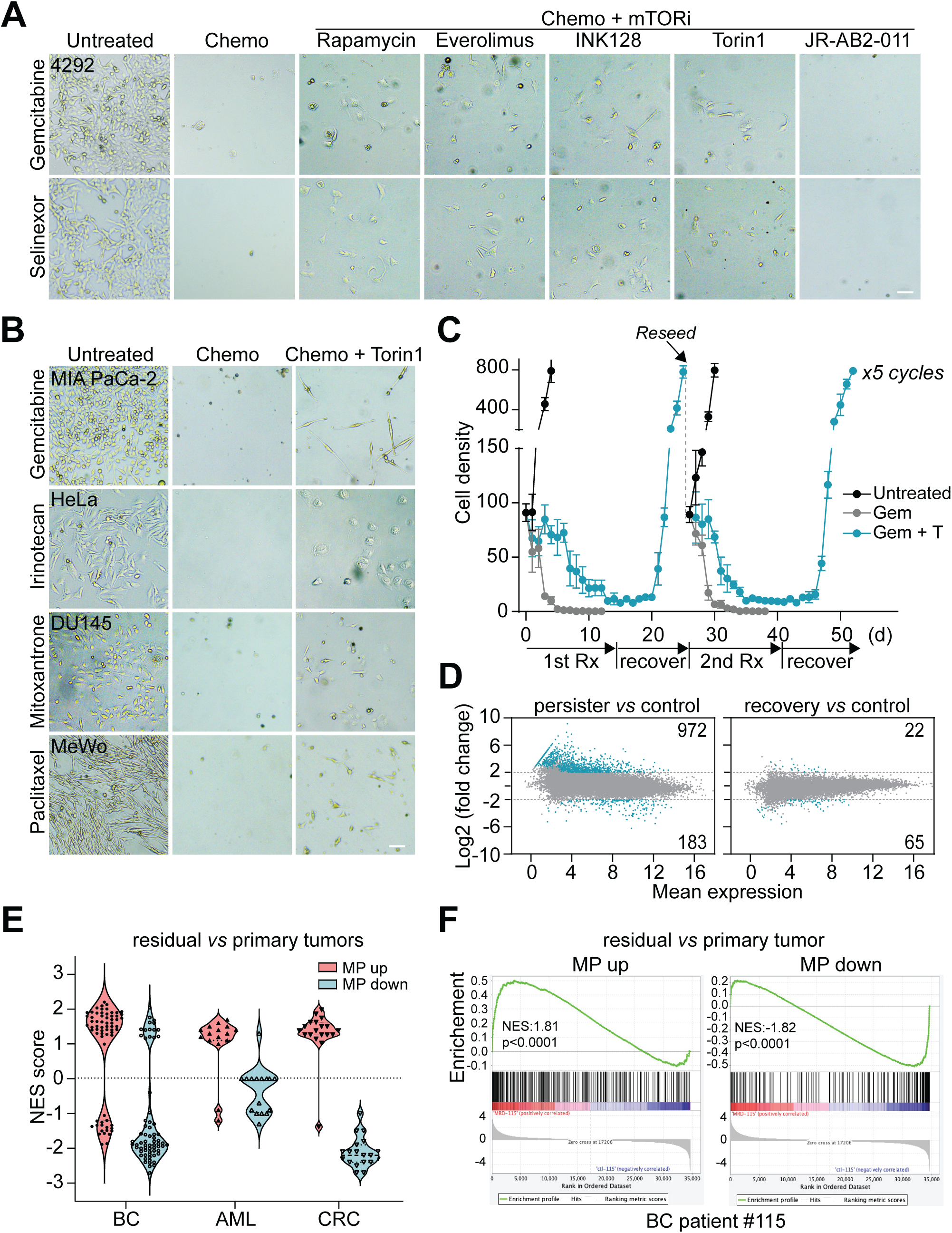
mTOR inhibition induces reversibly chemoresistant persisters. **A.** 4292 cells were treated with chemotherapeutics (2μM gemcitabine, 2μM selinexor), or chemotherapeutics plus a panel of mTOR inhibitors (100nM rapamycin, 100nM everolimus, 100nM Torin1, 100nM INK128, and 1μM JR-AB2-011) for seven days. Representative images of cell survival and morphology are shown. Scale bar, 100μm. **B.** Microscopic images of human cancer cells treated with Torin1 plus chemotherapeutic agents (MIA PaCa-2: 100nM gemcitabine; HeLa: 60μM irinotecan; DU145: 100nM mitoxantrone, and MeWo: 10nM paclitaxel). Scale bar, 100μm. **C.** Reversibility of chemoresistance in persister cells. Treatment (Rx), recovery, and reseeding process was repeated for five cycles with similar results. Y axis represents cell number in 50X field (0.64 mm^2^ of culture area). At least three independents fields were counted for each data point and cell numbers were plotted as mean ± SEM. **D.** Transcriptomic profiling of the control, persisters, and recovered cell populations. Black dots represent differentially expressed genes that change at least 4-fold (adjusted p < 0.001). **E.** Violin plots showing the GSEA results of residual *vs* primary tumors in multiple human cancer types using the MP signature. BC: breast cancer (GSE87455), AML: acute myeloid leukemia (GSE40442), CRC: colorectal cancer (GSE108277). NES: normalized enrichment scores. **F.** Representative GSEA plot of patient #115 from the breast cancer dataset (GSE87455).

To determine whether the induction of persisters following mTOR inhibition is a general phenomenon, we treated human pancreatic cancer MIA PaCa-2 cells with a panel of diverse chemotherapeutic agents. mTOR inhibition led to the emergence of persisters in the majority of agents tested, including gemcitabine (antimetabolite), paclitaxel (antimicrotubule), irinotecan (topoisomerase I inhibitor), mitoxantrone (anthracycline), doxorubicin (anthracycline), and etoposide (topoisomerase II inhibitor) (**Figure S2A**). The effects of mTOR inhibition were further evaluated in an additional panel of 30 human cancer cell lines from diverse tissue origins, including pancreatic, breast, prostate, colon, liver, lung, ovarian, cervical cancers and melanoma. Treatment with chemotherapeutic agents and mTOR inhibitors led to the emergence of persisters in the majority of these cells. Cell lines that failed to show the drug tolerant subpopulation were mostly TP53 wild type (MCF7, HCT116, H460, A549, A375, and HepG2), indicating that induction of the drug-tolerant state by mTOR inhibition may be a common feature of human cancers with TP53 mutations (**Figures 2B**, **S2B-C**).

We focused on the MIA PaCa-2 model to further examine the drug tolerant phenotype. Following drug removal, persister cells resumed proliferation. Recovered cells were sensitive to gemcitabine and completely eradicated by gemcitabine treatment alone, while combined treatment with gemcitabine and Torin1 again led to the emergence of survivors. The treatment-recovery was continued for five cycles and persisters consistently appeared following each cycle of chemotherapy plus mTOR inhibition (**Figure 2C**). The persister phenotype was further validated in three independent single-cell clones derived from MIA PaCa-2 cells, and induction of persisters and phenotypic reversibility was similarly observed in each clone (**Figure S3A**). These results indicate that the drug-tolerant cells arise *de novo* upon non-genetic adaptation and are unlikely to derive from rare clones with preexisting genetic mutations. Additionally, we compared the transcriptomes of control, persisters, and recovered cells (persisters that were cultured in drug free media for an additional 11 days). Persister cells displayed 972 upregulated and 183 downregulated genes (with at least 4-fold change and an adjusted p value of less than 0.001). Interestingly, the transcriptional changes are highly enriched for MYC target gene expression, G2/M cell cycle, mitotic transition, and mTOR signaling (**Figures 2D, S3B; Tables S2A-B**). While control and persisters differ drastically in their gene expression pattern, only modest differences are observed between the control and recovered cells. The top 200 most variable genes in persisters did not show significant differential expression in the recovered *vs* control cells (**Figure S3C)**. These results suggest that persisters largely reverts to the cellular state of the pre-treatment control following drug removal. The acquired resistance following mTOR inhibition is therefore unlikely mediated by stable genetic changes, but a consequence of an adaptive response to chemotherapeutic stress.

To examine the relevance of mTOR inhibition-induced persister phenotype in clinical cancer treatments, we derived an mTOR-regulated persister (MP) signature by identifying the common transcriptomic changes in the Torin1-treated *vs* control and persister *vs* control. The MP signature with 403 upregulated (MP up) and 213 downregulated genes (MP down) was used in Gene Set Enrichment Analysis (GSEA) of public transcriptomes of residual tumors following chemotherapy from several cancer types, including breast cancer^18^, colorectal cancer^19^, and acute myeloid leukemia^20^. Significantly, the upregulated MP genes were enriched, while the downregulated MP genes were depleted in the transcriptomes of residual tumors (**Figures 2E-F; Tables S3A-B**), suggesting that the mTOR-regulated persister state may be responsible for the minimal residual disease phenotype following clinical cancer chemotherapy.

### mTOR Activation increases chemosensitivity

Following the demonstration that mTOR inhibition promotes the development of drug-tolerant persisters, we examined whether activation of mTOR pathway could conversely increase chemosensitivity. We used CRISPR to knock out two genetic suppressors of mTOR, *TSC1* and *TSC2*, in MIA PaCa-2 cells. Western analysis confirmed the loss of *TSC1* or *TSC2* expression and activation of the mTORC1 pathway as indicated by increased S6 phosphorylation (**Figure 3A**). Single clones of the knockouts showed increased sensitivity to gemcitabine, 5-fluorouracil, selinexor, and paclitaxel, compared to the wild type clones (**Figure S4A**). We further labeled the wild type clones with red fluorescent protein (RFP) and *TSC1*/*TSC2* knockout clones with green fluorescent protein (GFP), and mixed at a one to one ratio in a multicolor competition assay (MCA). Treatment with gemcitabine or selinexor selectively eradicated the knockout clones as shown by disappearance of GFP positive cells, while the GFP/RFP ratio did not change significantly in the untreated populations (**Figures 3B-C**).

**Figure 3.**
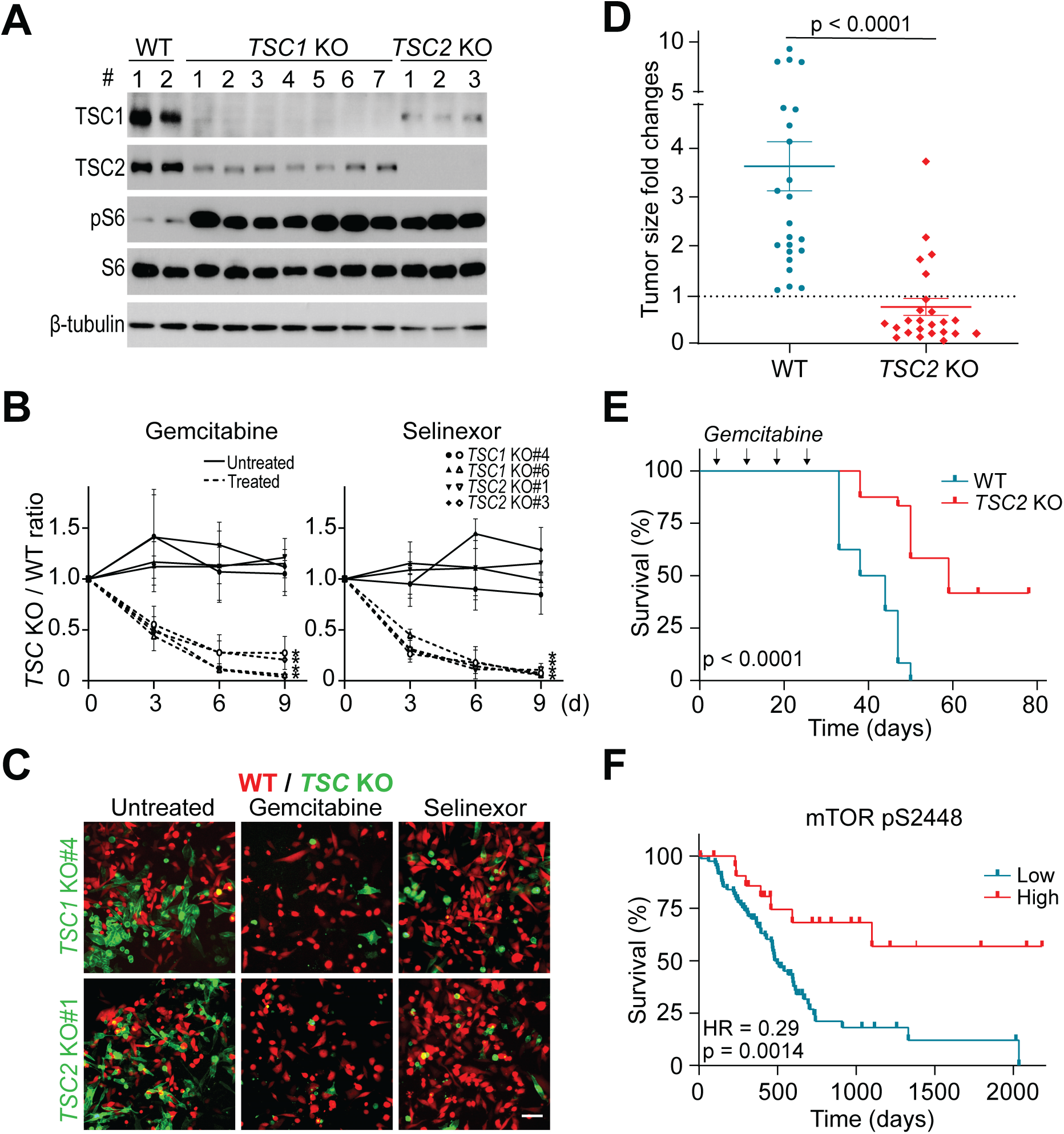
mTOR activation confers chemosensitivity *in vitro* and *in vivo*. **A.** Western analysis of single cell clones of *TSC1* and *TSC2* knockouts derived from MIA PaCa-2 cells. Two wild type (WT), seven *TSC1* knockout (*TSC1* KO), and three *TSC2* knockout (*TSC2* KO) clones were generated. TSC1, TSC2, phospho-S6 (pS6), and S6 proteins were detected to show genomic deletion and mTOR activation. β-tubulin serves as the loading control. **B.** MCA of wild type mixed with *TSC1* or *TSC2* knockout clones in the presence or absence of 30nM gemcitabine or 300nM selinexor for three, six, and nine days. Data were plotted as mean ± SEM. Two-tailed Student’s T test, * p < 0.001. **C.** Representative images of the MCA six days after the drug treatment. Scale bar, 100μm. **D.** Fold changes in tumor size based on BLI quantification following gemcitabine treatment in wild type and *TSC2* knockout tumors grown in NSG hosts. Two-tailed Student’s T test. **E.** Survival of NSG mice bearing wild type and *TSC2* knockout tumors following four weeks of gemcitabine treatment. Log-rank (Mantel-Cox) test, p < 0.0001. N=24 for wild type and N=25 for *TSC2* knockout xenografts. **F.** Survival analysis of human pancreatic cancer patients based on RPPA of phospho-mTOR-S2448 in tumors. Data from the TCGA study were analyzed using the TRGAted application.

To assess the chemosensitivity *in vivo*, we transplanted wild type and *TSC2^-/-^* MIA PaCa-2 cells into immunodeficient NOD *scid* gamma (NSG) mice. Despite four weeks of gemcitabine treatment, wild type tumors expanded four-fold according to bioluminescence imaging (BLI) quantification. Interestingly, significant shrinkage was observed in the majority of *TSC2^-/-^* tumors (**Figure 3D**). The increased chemosensitivity upon mTOR activation translated into better survival of the NSG hosts bearing *TSC2^-/-^* tumors (**Figure 3E**). Furthermore, analysis of the reverse phase protein array (RPPA) data in The Cancer Genome Atlas (TCGA) project^21^ points to a significant survival benefit in pancreatic cancer patients with high mTOR protein expression in their tumors (hazard ratio=0.381, p=0.022). In particular, high levels of phospho-mTOR-S2448, indicative of mTOR kinase activity, is remarkably correlated with better overall survival (hazard ratio = 0.29, p = 0.0014) and progress-free survival (hazard ratio = 0.414, p = 0.00049) in pancreatic cancer, as well as multiple other cancer types, including lung, liver, kidney, prostate, cervical cancers, and melanoma (**Figures 3F, S4B**). Taken together, mTOR inhibition promotes drug tolerance, while activation of the mTOR pathway increases chemosensitivity, and is predictive of improved clinical survival in cancer patients.

### Persister cells display a senescence phenotype

Persisters have a distinctly enlarged and flattened morphology, reminiscent of the human fibroblasts undergoing replicative senescence^22^. We therefore examined these cells for the molecular changes and phenotypic features associated with senescence. Flow cytometric analysis detected approximately 4- and 6-fold increase in the forward scatter (FSC) and side scatter (SSC) in the persisters compared to control, consistent with the characteristic cell size and granularity of senescent cells (**Figure S5A**). Torin1-treated cells demonstrated modest increase in FSC and SSC compared to the non-treated control. Canonical senescence-associated β-galactosidase (SABG) assay revealed extensive staining in the persisters but not in control or Torin1-treated cells (**Figures 4A-B, S5B-C**). Flow cytometric quantification of SABG activity with the 5-dodecanoylaminofluorescein di-β-D-galactopyranoside (C12FDG) substrate^23^ showed a 4-fold higher mean fluorescence intensity (MFI) in persister cells, while Torin1-treated cells did not differ from the control (**Figures 4C-D**). The senescent phenotype was also confirmed by drastic induction of p21 and γH2AX and silencing of lamin B1 protein expression in persisters as shown by Western analysis (**Figure 4E**). These markers remained unchanged in Torin1-treated cells, indicating the absence of senescence induction by mTOR inhibition. Additionally, persister cells displayed a typical senescence-associated secretory phenotype (SASP) as revealed by Luminex detection of secreted cytokines. A minimum of 4-fold induction by the combination treatments was observed in multiple cytokines commonly associated with SASP, including CCL2, CCL26, G-CSF, MIF, FGF2, and CXCL5 (**Figure 4F**)^24–27^. Finally, analysis of our transcriptomic data sets utilizing a senescence classifier with a 97% specificity^28^, clearly identified the persisters as the senescent population (score of 0.99), while control, Torin1-treated, and recovered cells did not show any sign of senescence (scores less than 0.01) (**Figure 4G**). Taken together, drug-tolerant persister cells display a prominent senescence phenotype according to multiple independent measures. Importantly, persister cells recover and re-establish the tumor cell population following drug removal with complete reversibility, demonstrating a cellular state^29, 30^ distinctly different from the replicative senescence with irreversible cell cycle arrest.

**Figure 4.**
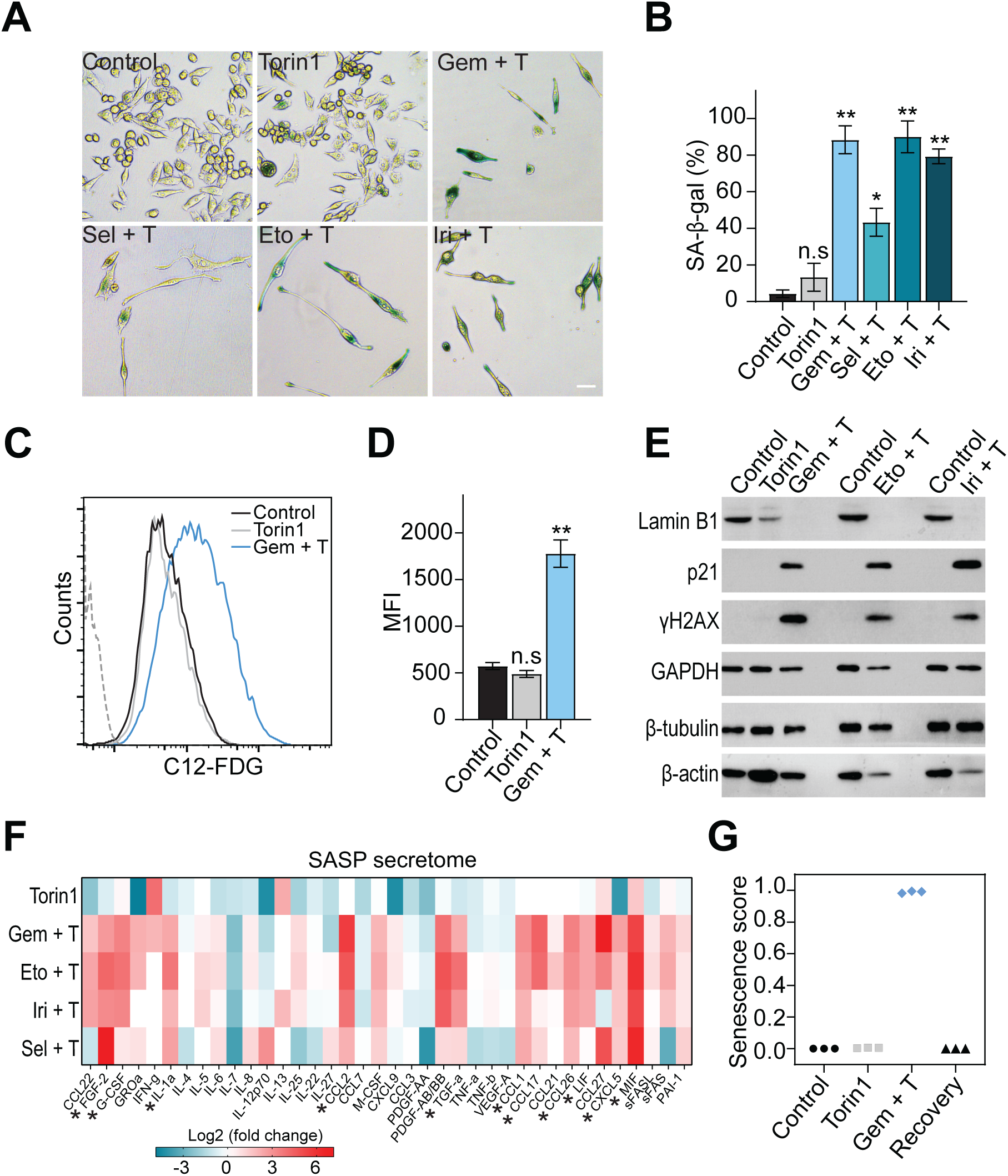
Persister cells display a senescence phenotype. **A.** SABG staining of MIA PaCa-2 cells treated with 100nM Torin1, or Torin1 plus chemotherapeutic agents (100nM gemcitabine, 2μM selinexor, 6μM etoposide, and 6μM irinotecan) for eight days. **B.** Quantification of SABG positive cells in control, Torin1, or Torin1 plus chemotherapeutic agents. Student’s T test. ** p < 0.001; * p < 0.01; n.s: not significant. **C-D.** Flow cytometric quantification of SABG activity using the C12-FDG fluorescent substrate in MIA PaCa-2 cells treated with 100nM Torin1 or 100nM gemcitabine plus Torin1 for nine days. Grey dashed line indicates the unstained control. Student’s T test. ** p < 0.001; n.s: not significant. **E.** Western analysis of senescence markers in cells treated with Torin1 or Torin1 plus chemotherapy for nine days. **F.** Heatmap of cytokine secretion in MIA PaCa-2 cells following a nine-day treatment with Torin1, or Torin1 plus chemotherapeutic (100nM gemcitabine, 1μM selinexor, 6μM etoposide, and 6μM irinotecan). Cytokines were detected using the Luminex assays. Asterisks indicate senescence-associated cytokines upregulated in the persisters. **G.** Senescence scores for control, Torin1-treated, persisters, and recovered cells analyzed using a senescence classifier from the Cancer SENESCopedia. Each condition contains three biological replicates of RNAseq data.

### A small-molecule chemical library screen reveals survival mechanisms of persisters

To identify the survival mechanisms in persisters, we performed a chemical library screen in the persister model of MIA PaCa-2 cells treated with gemcitabine and Torin1 for nine days, using a small-molecule library with approximately 200 inhibitors for various druggable targets and cellular processes (**Table S4A**). Each inhibitor was tested in a serial dilution in both parental and persister cells. While most inhibitors did not affect persister survival, a few compounds targeting G2/M checkpoints, autophagy, and cell death pathways (apoptosis and ferroptosis^31^) selectively eradicated persisters, with minimal effect on the parental cells (**Figures 5A-B, S6; Tables S4A-B**).

**Figure 5.**
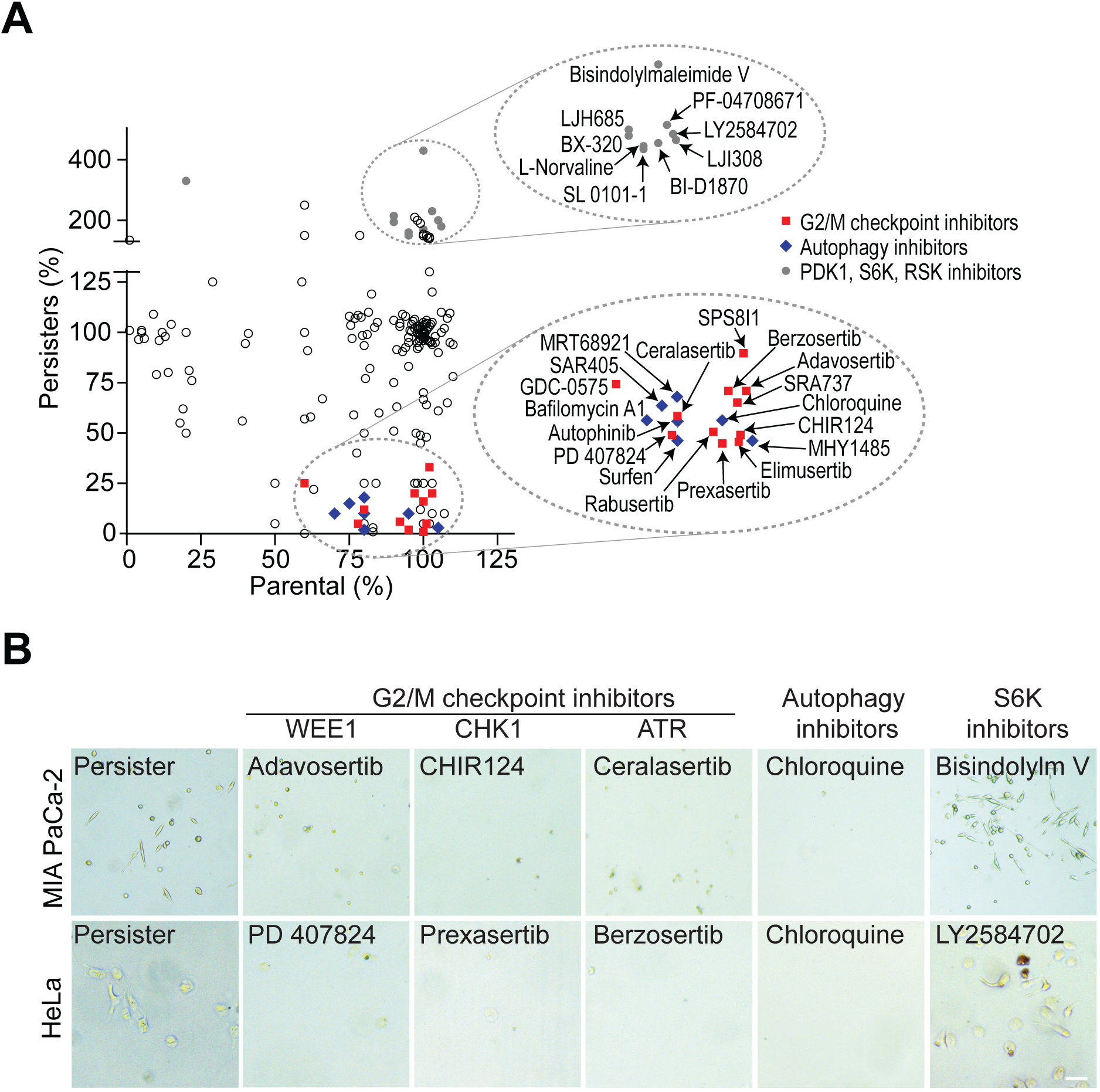
Multifaceted survival mechanisms of persisters revealed by a small molecule chemical screen. **A.** The effects of inhibitors on survival of persisters and parental cells. Each chemical was plotted based on its activity in parental and persister cells. For each drug, data were normalized to untreated parental and persister cells. Circled dots signify three major groups of drugs that reduce or promote survival of persisters. **B.** Representative images of persister survival following treatment with inhibitors from the chemical library. Persister cells were induced by combined treatment with 100nM gemcitabine and 100nM Torin1 for MIA PaCa-2 cells, and 60μM irinotecan and 100nM Torin1 for HeLa cells. Representative inhibitors targeting G2/M checkpoint, autophagy, and S6 phosphorylation are shown, including 100nM adavosertib, 100nM CHIR124, 10μM ceralasertib, 10μM chloroquine, and 30μM bisindolylmaleimide V for MIA PaCa-2 cells and 100nM adavosertib, 1μM prexasertib HCI, 3μM PD 407824, 10μM chloroquine, and 1μM LY2584702 for HeLa cells.

Persister cells were dependent on autophagy for survival, as general inhibitors of autophagy, such as chloroquine and bafilomycin, and specific inhibitors, including SAR405 (VPS34 inhibitor) and MRT68921 (ULK1/2 inhibitor), efficiently eradicated these cells (**Figures 5A-B**). In parallel, we observed strong evidence of autophagy induction in persisters based on Western analysis of protein markers (phospho-CREB, phospho-ATG14, and LC3II), detection of LC3 puncta using a dual-color DsRed-LC3-GFP fluorescent reporter^32^, and GSEA of transcriptomes (**Figures S7A-E**). Sensitivity to autophagy inhibitors was further confirmed in MIA PaCa-2 persister cells induced by treatment with selinexor, etoposide, irinotecan, and doxorubicin plus mTOR inhibitors, as well as additional cancer types and chemotherapeutic contexts (**Figures S6A, S7F-G**). Addition of chemical inhibitors at the beginning of the treatment or following the appearance of persisters, equally blocked their survival (**Figure S6B**). Furthermore, inhibition of multiple key G2/M cell cycle checkpoint regulators, including CHK1, ATR, and WEE1, eliminated the persisters (**Figure 5B**), demonstrating that activation of G2/M cell cycle checkpoint is essential for persister survival. Conversely, multiple chemicals targeting CDK4/6, PDGFRs, PDK1, RSKs, and S6Ks, increased the persister population (**Figures 5A-B, S6C; Tables S4A-B**). In particular, a few kinases, such as PDK1, RSKs, and S6Ks, functionally converge on the S6 phosphorylation downstream of mTORC1 pathway, suggesting that emergence of persister cells following mTOR inhibition may be due to reduced S6 phosphorylation. Thus, our chemical screen identifies the multifaceted survival mechanisms in persisters.

### mTOR suppression potentiates G2/M cell cycle arrest and promotes survival of persisters

Our chemical screen identified multiple G2/M checkpoint inhibitors targeting CHK1, ATR, and WEE1, effectively eliminating the persisters, which were further confirmed in multiple chemotherapeutic contexts (**Figures 5A-B, S8A**). In a recent study, acute myeloid leukemia cells treated with chemotherapy persisted through a senescence-like state, which was effectively targeted by ATR inhibitors^30^. We, therefore, assessed cell cycle distribution of the persisters to unravel the survival mechanisms involving G2/M regulation. Flow cytometric analysis of DNA copy number in persisters induced by multiple drug combinations showed prominent G2/M cell cycle arrest and polyploidy (**Figure S8B**). Following release from the combined treatments, persisters recovered normal proliferation and displayed cell cycle profiles comparable to that of control cells. Using a fluorescent ubiquitination-based cell cycle indicator (FUCCI) reporter, we showed that treatment with chemotherapy plus Torin1 resulted in G2/M cell cycle arrest, while Torin1-treated cells were predominantly in G1/S phase (**Figures 6A-B**). Additionally, induction of phospho-CHK1 (Ser345) and phospho-WEE1 (Ser642) pointed to activation of G2/M checkpoint in persisters (**Figures 6C-D**). In stark contrast to the G2/M checkpoint activation, expression of G2/M transition genes was drastically downregulated in persister cells. Western analysis showed significant reduction in PLK1 and cyclin B1 protein levels in persisters, similar to Torin1-treated cells (**Figures 6C-D**). Furthermore, GSEA of our transcriptomic data revealed depletion of transcripts involved in G2/M transition in both Torin1-treated and persister cells (**Figures 6E-F**). Thus, although persister cells are predominantly accumulated at G2/M phase, they are unable to progress through cell cycle due to significant downregulation of genes involved in G2/M transition by mTOR inhibition, thereby avoiding mitotic catastrophe caused by premature mitotic entry. Concordantly, in a previous study, mTOR inhibition potentiated cell cycle arrest after irradiation through the transcriptional downregulation of cyclin B1 and PLK1, leading to increased survival of cancer cells following DNA damage^33^. Thus, regulation of the G2/M cell cycle arrest by mTOR inhibition mechanistically contributes to the survival of persister cells.

**Figure 6.**
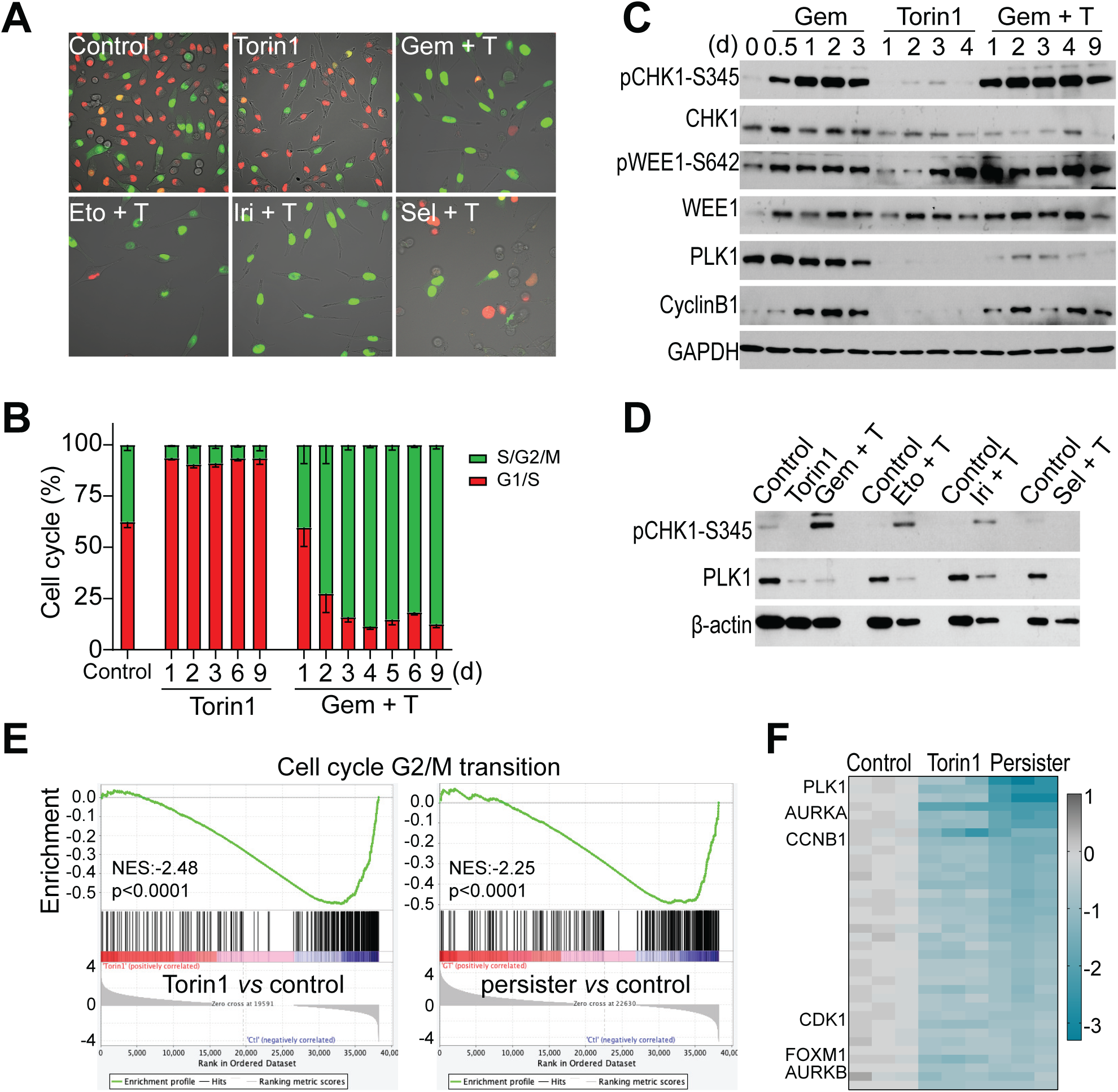
mTOR inhibition potentiates G2/M cell cycle arrest and promotes persister survival. **A-B.** Cell cycle tracking with FUCCI reporter in control, Torin1-treated, and persister cells. Scale bar, 100μm. **C.** Western analysis of key proteins involved in G2/M checkpoint activation and G2/M cell cycle progression. MIA PaCa-2 cells were treated with 100nM gemcitabine, 100nM Torin1, or the combination. **D.** Western analysis of phospho-CHK1 and PLK1 in MIA PaCa-2 cells treated with Torin1 or Torin1 plus chemotherapeutic agents (100nM gemcitabine, 2μM selinexor, 6μM etoposide, and 6μM irinotecan). **E.** GSEA of G2/M transition signature (GOBP_CELL_CYCLE_G2_M_PHASE_TRANSITION from MSigDB) in Torin1-treated and persister cells. Normalized enrichment scores (NES) and nominal p values are provided. **F.** Expression of G2/M phase specific transcripts in the control, Torin1-treated, and persister cells. Genes were derived from the GSEA G2/M transition signature in Figure 6E. Key regulators of G2/M transition are labeled. The color bar indicates the scale of log2 fold changes.

## DISCUSSION

We have identified mTOR pathway as a key regulator of chemosensitivity through CRISPR screening. Chemotherapeutic treatments in the presence of mTOR inhibition induce a drug-tolerant subpopulation of tumor cells with a senescence phenotype, the survival of which is dependent on autophagy and G2/M cell cycle arrest. Conversely, activation of mTOR pathway confers increased chemosensitivity in cancer cells with genetic ablation of *TSC1/2*. Additionally, activated mTOR signaling as indicated by phospho-mTOR-S2448 levels in patients’ tumors predicts better clinical survival among multiple cancer types. This notion is further supported by reports of increased sensitivity to DNA damage-inducing therapies in cancer patients with mTOR activation caused by germline and/or somatic mutations of *TSC1/2*^34–37^. These observations point to a heightened chemosensitivity due to mTOR activation and highlight the tumor suppressor activity of mTOR in the context of cancer treatments, despite its well-characterized oncogenic functions in tumorigenesis.

Persisters display a senescence phenotype, distinct from replicative senescence with irreversible arrest^38^, and similar to the physiological diapause. mTOR is shown to causally regulate diapause during mouse blastocyst development^39^. More recently, colorectal cancer cells reportedly enter a diapause-like drug-tolerant persister state, concomitant with the reduction of mTOR signaling^5^. The functional roles of mTOR in physiological diapause upon nutritional deprivation and chemotherapy-induced senescence are likely similar, both involving induction of autophagy and cell cycle arrest^39^. Under physiological diapause, mTOR inhibition induces autophagy and dormancy to support long-term survival and pluripotency of blastocysts^39^. Under the stress of chemotherapy, mTOR inhibition promotes cancer cells to adopt a reversible drug-tolerant senescence state, which supports survival and allows for effective DNA repair, but incidentally, leading to chemoresistance and tumor recurrence.

While mTORC1 inhibitors, such as everolimus and temsirolimus, are approved for treatment of several cancer types, including pancreatic neuroendocrine tumor, breast cancer, and renal cell carcinoma, the clinical applications are focused on controlling tumor expansion in late-stage cancers. We observe that mTOR suppression promotes survival of cancer cells undergoing chemotherapy and dampens therapeutic efficacy. Similarly, chemoresistance in colorectal cancers is reportedly associated with reduced mTOR activity^5^. Thus, the mTOR pathway may play a tumor suppressing role in the context of chemotherapy. This may explain the lack of efficacy of mTOR inhibitors in cancer clinical trials and argues against their widespread use in combination with DNA damage-inducing agents. In this light, stimulation of mTOR signaling, but not its inhibition, may be tested to reduce residual diseases and improve chemotherapeutic efficacy.

The paradoxical roles of oncogenes in regulating chemotherapeutic sensitivity is not limited to mTOR. Overexpression of MYC oncogene, while inducing aggressive tumor behavior, in fact leads to increased chemosensitivity^40–42^. Additionally, silencing MYC induces a diapause-like cellular state that is chemoresistant^43^. Not surprisingly, MYC is one of the top enriched hits in our CRISPR screen (**Figure 1B**). Tumor suppressor activity may confer chemoresistance. Disruption of the p53 tumor suppressor enhances sensitivity to multiple chemotherapeutic agents, while its expression results in chemoresistance in various cancer types^44–47^. Similarly, attenuation of the RB tumor suppressor leads to aggressive tumorigenic growth accompanied by increased chemosensitivity^48–50^. Regulation of chemoresistance by oncogenes and tumor suppressors may mechanistically converge on senescence as MYC inhibition and p53/RB activation are known inducers of senescence^44, 51–56^. Depending on specific cellular contexts, suppression of oncogenes or activation of tumor suppressors may not result in tumor eradication via various forms of cell death, but instead lead to tumor persistence through a reversible senescence state, thus promoting tumor recurrence. The seemingly paradoxical roles of oncogenes and tumor suppressors in regulating tumor growth *vs* therapeutic sensitivity may be a common theme.

## METHODS

### Cell lines and chemical reagents

The 4292 murine PDAC cell line was a gift from Dr. Marina Pasca di Magliano^12^. All human tumor cell lines were from ATCC. Cells were cultured in DMEM media (except for T47D, MDA-MB-231, C4-2B, DU145, H460, and 22Rv1, which were cultured in RPMI1640, and PC3 in F-12K), supplemented with 10% fetal bovine serum and 1% penicillin/streptomycin in a 5% CO2 humidified incubator. Gemcitabine, 5-fluorouracil, paclitaxel, carboplatin, Torin1, rapamycin, everolimus, temsirolimus, bafilomycin, chloroquine, MHY1485 were purchased from Cayman Chemical. Selinexor, SAR405, and MRT68921 were obtained from Selleckchem. All chemicals for the small-molecule library screening were purchased from Cayman Chemical.

### CRISPR Cas9 genome-wide library screening

CRISPR library screen was performed in biological replicates following a published protocol^57^. 4292 cells were infected with lentiCas9-Blast (Addgene #52962). Following blasticidin selection, cells were infected at low viral titer with the pooled mouse CRISPR lentiviral library containing 78,637 gRNAs targeting 19,674 genes (Addgene #73633-LV). Infected cells were selected with puromycin (1 μg/ml) for two days. The experimental pools were treated with gemcitabine (20nM) or selinexor (0.33μM) for 12 days. All cell pools were passaged every three days with at least 50 million cells per pool to ensure a minimum of 500X coverage. Following drug treatment, sgRNA libraries in the cell populations were isolated by PCR amplification and identified by Hiseq. Computational analysis of the sgRNA libraries was performed using MAGeCK (version 0.5.9.2) as reported^15^.

### CRISPR knockout

The guide RNAs targeting specific genes were designed using CHOPCHOP (https://chopchop.cbu.uib.no/). Two sgRNAs were cloned in pSpCas9(BB)-2A-GFP (PX458) (Addgene, #48138). MIA PaCa-2 cells were transiently transfected with an equimolar mixture of the sgRNAs using lipofectamine 3000 (Thermo Fisher). Three days after transfection, GFP positive cells were sorted for single cells on a BD FACS Aria cell sorter (BD Biosciences). Gene knockout was confirmed by genomic DNA PCR and Western analysis. The sgRNA sequences are:

hTSC1-A CGAGATAGACTTCCGCCACG
hTSC1-B AGTCGGTGGGAGACGACTAT
hTSC1-C GACGTCGTTGTCCTCACAAC
hTSC1-D TACCAATGATTCCACAGTCT
hTSC2-A CGTCTGCGACTACATGTACG
hTSC2-B AGGAGACGACTCGCTCGATG
hTSC2-C CGTCCGGACCGCGTCCTCTG
hTSC2-D CTGTCGCACCATCAACGTCA.

### SABG staining

MIA PaCa-2 and MDA-Panc-28 cells were plated in 24-well plates at 10,000 cells/well and treated with chemotherapeutic agents and Torin1 the next day. Following eight days of drug treatment, cells were stained using Senescence Cells Histochemical Staining Kit (Sigma #CS0030) according to the manufacturer’s protocol. Nine images from each well were acquired and positively stained cells were manually counted.

### Luminex assays of secreted cytokines

Secretome analysis was performed using Luminex Human 80 plex (EMD-Millipore) at Stanford Human Immune Monitoring Center. Cell culture media were changed three days before sample collection. For each sample, levels of cytokines were normalized to the media volume and cell number. All cytokine detection was performed in technical duplicates.

### RNAseq analysis

MIA PaCa-2 untreated cells (control), treated with 100nM Torin1 (Torin1-treated), or 100nM Torin1 and 100nM gemcitabine (persisters) for nine days, and persister cells that were grown in drug free media for 11 days (recovery) were used for RNAseq analysis. Total RNA was extracted using Qiagen RNeasy Plus kit (Qiagen). Library construction and total RNA sequencing were performed commercially (Novogene). For RNA-seq data analysis, kallisto (version 0.46.2)^58^ was used to pseudoalign paired-end reads to a reference transcriptome with 100 bootstraps. The reference transcriptome used to build the kallisto index consisted of the “Protein-coding transcript sequences” and “Long non-coding RNA transcript sequences” FASTA files from Human Release 38 (GRCh38.p13) of GENCODE. The corresponding GENCODE human primary assembly GTF file was used as an annotation file. The R library, sleuth (version 0.30.0)^59^, was used to produce normalized gene-level abundance estimates and perform differential gene expression analysis.

### GSEA using the mTOR-regulated persister signature

To generate the mTOR-regulated persister signature from the RNAseq data, we first derived the genes differentially expressed in Torin1-treated and persister cells (two-fold changes at an adjusted p<0.01). The overlap of the two sets of differentially expressed genes was defined as the mTOR-regulated persister signature, which includes both upregulated and downregulated genes (MP up and MP down) (**Table S3A**). GSEA of the public datasets (GSE87455 for breast cancer, GSE108277 for colorectal cancer, and GSE40442 for acute myeloid leukemia) were performed using our MP signature on post-chemotherapy residual tumors *vs* pre-treatment primary tumors. The normalized enrichment scores (NES) for both MP up and MP down signatures in the three public data sets are plotted using GraphPad Prism (version 8.4.2).

### Cell cycle analysis using FUCCI reporter

FUCCI reporter (Addgene #86849) was stably expressed in MIA PaCa-2 cells by lentiviral transduction. Cells were seeded in 24 well-plates and treated the next day with indicated drugs for nine days. At least nine representative images of each condition were captured on a FV3000 confocal microscope. Red and green fluorescence indicate distribution in the G1/S and S/G2/M phase, respectively.

### Survival analysis of patients’ RPPA data in TCGA

Survival analysis of patients’ samples were based on the reverse-phase protein array (RPPA) data on mTOR and phospho-mTOR-S2448 in The Cancer Genome Atlas (TCGA) project. Data analysis was performed through the TRGAted application using optimal cutoff^21^.

### Western blot analysis

Cells were lysed on ice in CelLyticTM MT reagent (Sigma) supplemented with protease and phosphatase inhibitors. Protein concentration was determined with BCA Protein Assay Kit (Thermo Fisher). Protein lysates were resolved on SDS-PAGE gels and routine procedures were followed. Primary antibodies include TSC1 (#6935), TSC2 (#4308), phospho-S6 (#4858), S6 (#2217), p21 (#2947), Lamin B1 (#12586), LC3 A/B (#12741), phospho-ATG14-s29 (#92340), phospho-CREB-S133 (#9198), phospho-CHK1-S345 (#2348), phospho-WEE1 (#4910), WEE1(#13084), PLK1 (#4513), γH2A.X (#9718) from Cell signaling, β-tubulin (#10094-1-ap), β-actin (#20536-1-ap), and GAPDH (1:10000, #10491-1-ap) from ProteinTech, and CHK1 (#sc- 8408) and Cyclin B1(#sc-245) antibodies from Santa Cruz.

### Flow cytometry analysis

For cell cycle distribution, cells were resuspended in PBS, fixed with ice-cold ethanol, and treated with RNase and propidium iodide and analyzed on a BD FACS Fortessa flow cytometer (BD Bioscience). Quantification of SABG enzymatic activity followed a previously described protocol^60^. Briefly, cells were treated with bafilomycin A1 (100nM) for one hour and stained with C12-FDG (33μM) for two hours at 37℃ under 5% CO2 before flow cytometric analysis. Data were analyzed using FlowJo software (Tree Star).

### Autophagic reporter assay

pQCXI Puro DsRed-LC3-GFP (Addgene #31182) reporter was introduced into MIA PaCa-2 cells by retroviral infection. Cells expressing the reporter were seeded into 4-well chambered coverslip (ibidi #80426) and treated with the indicated drugs for nine days. Images were acquired using a FV3000 confocal microscope. For each condition, LC3 puncta in at least 50 cells were counted.

### Cell viability assay

Two thousand MIA PaCa-2 wild type or *TSC1*/*TSC2* knockout cells were seeded in 96-well plates. Drugs were added the next day in four replicates. Cell viability was detected using the CellTiter-Glo 2.0 Cell Viability Assay (Promega) following four to five days of drug exposure. Calculation of the half-maximal inhibitory concentration (IC50) was carried out with GraphPad Prism (version 8.4.2).

### Multicolor competition assay

MIA PaCa-2 wild type (WT) and *TSC1*/*TSC2* knockout (KO) cells were transduced with retrovirus expressing RFP (pQCXIP-turboRFP, Addgene #73016) or EGFP (pQCXIP-EGFP-F, Addgene #73014). MIA PaCa-2 WT-RFP were mixed 1:1 with *TSC1*/*TSC2* KO-GFP cells. One day after seeding (d0) and 3, 6, 9 days after chemotherapy (d3, d6, d9), cells were imaged on a Leica DMi8 fluorescence microscope with a 10X objective and RFP and GFP positive cells were counted. Relative cell fitness was calculated from the ratio of RFP/GFP positive cells normalized to the d0 value.

### Small-molecule chemical library screen

Approximately 200 chemical inhibitors covering major cell signaling and survival pathways were purchased from Cayman Chemical and dissolved in DMSO or PBS. Persister cells were induced by combined treatment of MIA PaCa-2 cells with gemcitabine and Torin1. Inhibitor was diluted 1:3 at a concentration range of 30nM-30µM, and added to the parental cells (single-agent treatment) and persister cells (triple treatment) for nine days. Cell survival were monitored daily under microscope. Persisters in the triple-treated wells were recorded on days six and nine, and compared to the untreated persisters induced by gemcitabine plus Torin1. For each inhibitor, single-agent treatment of the parental cells serves as a control to rule out non-specific toxic effects.

### In vivo therapeutic studies

Approximately four million firefly luciferase-labelled MIA PaCa-2 WT and *TSC2* KO cells were mixed with Matrigel and transplanted subcutaneously in NSG hosts. Tumors were allowed to grow for three weeks prior to gemcitabine treatment (100mg/kg, twice per week) for four weeks. Post-treatment survival was monitored for six weeks. BLI was performed on an IVIS Lumina III platform. Data were analyzed using the Living Image software (v4.2). All animal experiments were approved by IACUC at Houston Methodist Research Institute and were performed in accordance with institutional and national guidelines.

### Statistical analysis

Two-tailed Student’s t test was used to analyze the MCA data. Cell death rates among different treatment groups were analyzed using ANOVA with Tukey’s test. Results were presented as mean ± SEM. Animal survival was compared by Kaplan-Meier analysis with the log-rank (Mantel-Cox) test.

## Data availability

Sequencing data generated in this study have been deposited in GEO with accession number GSE162065 for the CRISPR screens and GSE189764 for the RNAseq experiment.

## Acknowledgements

We thank Dr. Marina Pasca di Magliano for generously providing the 4292 murine pancreatic cancer cell line, and Dr. Yael Rosenberg-Hasson for assistance with the cytokine assays. This work was supported in part by NIH K22CA207598 (Y.Li.), CPRIT RP200472 (Y.Li.), and NIH T32GM008042 (D.K.S.; UCLA-Caltech Medical Scientist Training Program).

## Author Contributions

Y.Liu performed *in vitro* studies and chemical screens, and analyzed the public data sets. N.G.A. carried out CRISPR screens and *in vivo* studies. D.K.S. analyzed the CRISPR screen and RNAseq data. Y.Li conceived and supervised the project. Y.Liu, N.G.A, and Y. Li wrote the manuscript.

## Supplementary Figure Legends

**Figure S1. Enrichment of positive regulators, and depletion of negative regulators of mTOR signaling in a CRISPR screen of DNA damaging agents.**

The CRISPR screen reported by Oshima K et al, (Nature Cancer. 2020. PMID: 33796864) was performed in REH acute lymphoblastic leukemia cells, which carry a TP53 mutation. The cells were treated with seven chemotherapeutic agents (VCR: vincristine; 6MP: 6-mercaptopurine; L-ASP: L-asparaginase; AraC: cytarabine; MTX: methotrexate; DNR: daunorubicin; and MAF: maphosphamide). Red and blue asterisks denote positive and negative regulators of the mTOR signaling pathway. The color bar indicates the scale of log2 fold changes.

**Figure S2. mTOR inhibition induces drug-tolerant persisters.**

**A.** Brightfield images of MIA PaCa-2 cells treated with chemotherapeutic agents (100nM doxorubicin, 2μM selinexor, 10nM paclitaxel, 6μM etoposide, 60nM mitoxantrone, and 6μM irinotecan), in the presence or absence of 100nM Torin1 for nine days. Scale bar, 100μm.

**B.** Representative images of persister cell recovery following drug withdrawal. Top panel: PC3 cells were treated with 300nM paclitaxel plus 100nM Torin1 for nine days (d0), followed by drug withdrawal for ten days (d10) and 60 days (d60). Middle panel: T-47D cells were treated with 40μM carboplatin plus 100nM Torin1 for nine days (d0), followed by drug withdrawal for ten days (d10) and 30 days (d30). Bottom panel: MeWo cells were treated with 10nM paclitaxel plus 30nM Torin1 for 12 days (d0), followed by drug withdrawal for ten days (d10) and 30 days (d30). Scale bar, 100μm.

**C.** Persister cell phenotype in a panel of 30 human cancer cell lines from diverse cancer types. Cells were treated with either chemotherapeutic agents or chemotherapy combined with Torin1. Cell lines with wild type TP53 were indicated. HNSCC: head and neck squamous cell carcinomas.

**Figure S3. The persister phenotype is reversible.**

**A.** Reversibility of persister phenotype in three independent single-cell clones derived from MIA PaCa-2 cells. Y axis represents cell number in 50X field (area of 0.64 mm^2^). Cells in at least three independents fields were counted for each data point and the cell numbers were plotted as mean ± SEM.

**B.** GSEA results of hallmark signatures from MSigDB in persister *vs* control, and Torin1-treated *vs* control comparisons.

**C.** Heatmap of the expression of 200 most variable genes from persister *vs* control comparison, in control, persister, and recovered cells. The color bar indicates the scale of log2 fold changes.

**Figure S4. Activation of mTOR increases chemosensitivity.**

**A.** Drug sensitivity of wild type and *TSC1* or *TSC2* knockout clones of MIA PaCa-2 cells to gemcitabine, selinexor, 5-fluorouracil (5-FU), and paclitaxel. Cells were treated with each drug for four to five days.

**B.** Survival analysis of various human cancer patients based on RPPA of phospho-mTOR-S2448. Data were derived from the TCGA study and analyzed using the TRGAted application. Y axis indicates the fraction of survival while X axis shows days of follow up. ACC: Adrenocortical carcinoma, CESC: Cervical squamous cell carcinoma and endocervical adenocarcinoma, KIRC: Kidney renal clear cell carcinoma, KIRP: Kidney renal papillary cell carcinoma, LIHC: Liver hepatocellular carcinoma, LUAD: Lung adenocarcinoma, MESO: Mesothelioma, PRAD: Prostate adenocarcinoma, SARC: Sarcoma.

**Figure S5. The senescence phenotype of persisters.**

**A.** Flow cytometric analysis of FSC and SSC in MIA PaCa-2 cells treated with Torin1 or Torin1 plus gemcitabine for nine days.

**B-C.** SABG staining and quantification of MDA-Panc-28 cells treated with Torin1, or Torin1 plus chemotherapeutic agents for six days (10μM gemcitabine, 1μM selinexor, 3μM etoposide, and 10μM irinotecan). Student’s T test. ** p < 0.001, n.s: not significant, p > 0.05.

**Figure S6. A small-molecule chemical library screen reveals survival mechanisms of persisters.**

**A.** Representative images of MIA PaCa-2 cells treated with various chemotherapeutics (2μM selinexor, 6μM etoposide, 6μM irinotecan and 100nM doxorubicin) plus 100nM Torin1, or in combination with 10μM chloroquine or 100nM adavosertib for nine days. Scale bar, 100μm.

**B.** Representative images of MIA PaCa-2 cells treated with 100nM gemcitabine and 100nM Torin1 (GT) (top panel), with 100nM adavosertib or 10μM chloroquine added at the beginning of drug treatment (GT + inhibitor) (middle panel), or five days after the start (GT to inhibitor) (bottom panel). Scale bar, 100μm.

**C.** Representative images of MIA PaCa-2 Cells treated with 100nM gemcitabine and 100nM Torin1, and autophagy inhibitors (2nM bafilomycin A1, 20μM MHY1485, 20μM ULK101), and G2/M checkpoint inhibitors (100nM berzosertib, 300nM SPS8I1, 3μM SRA737). CDK4/6 inhibitor (600nM abemaciclib), PDK1 inhibitor (3μM GSK2334470), RSK inhibitor (30μM LJI308). Scale bar, 100μm.

**Figure S7. High autophagy activity is essential for persister survival.**

**A.** LC3-II induction in MIA PaCa-2 and Panc-1 cells following treatment with 100nM Torin1 for nineteen days.

**B.** Western analysis of phospho-CREB, phospho-ATG14, and LC3-II in persister cells treated with Torin1 plus chemotherapeutic agents (100nM gemcitabine, 2μM selinexor, 6μM etoposide, and 6μM irinotecan).

**C-D.** Representative images and quantification of LC3 puncta in MIA PaCa-2 treated with Torin1 or Torin1 plus the chemotherapeutic agents. Student’s T test. ** p < 0.001, * p < 0.01. Scale bar, 100μm.

**E.** GSEA of autophagy signature (GOCC_AUTOPHAGOSOME_MEMBRANE from MSigDB) in Torin1-treated and persister cells. NES and nominal p values are shown.

**F.** Representative images of MIA PaCa-2 cells treated with 100nM gemcitabine plus 100nM Torin1 and autophagy inhibitors (10μM SAR405, 5μM MRT68921), and HeLa cells treated with 60μM irinotecan plus 100nM Torin1 and autophagy inhibitors (10μM SAR405, 3nM bafilomycin A1). Scale bar, 100μm.

**G.** Representative images of eradication of persister cells in MDA-MB-231, DU145, and SW480 following chloroquine treatment. Persisters were induced in MDA-MB-231 by treatment with 10nM paclitaxel and 100nM Torin1, DU145 by 10nM paclitaxel and 300nM Torin1, and SW480 by 1μM gemcitabine and 300nM Torin1. Scale bar, 100μm.

**Figure S8. G2/M checkpoint activation and cell cycle arrest in persisters.**

**A.** Representative images of eradication of persisters in MDA-MB-231, DU145, and SW480 following adavosertib treatment. Persisters were induced in MDA-MB-231 by treatment with 10nM paclitaxel and 100nM Torin1, DU145 by 10nM paclitaxel and 300nM Torin1, and SW480 by 1μM gemcitabine and 300nM Torin1. Scale bar, 100μm.

**B.** Cell cycle analysis by flow cytometry following propidium iodide (PI) staining in control, persister, and recovered cells. The percentage of cells in different phases of the cell cycle was labeled in the top right corner. MIA PaCa-2 cells were treated with Torin1 plus chemotherapeutic agents (100nM gemcitabine, 2μM selinexor, 6μM etoposide, and 6μM irinotecan) for nine days (persisters) and cultured in drug free media for an additional 9-12 days (recovered cells).

## Supplementary Tables

**Table S1.** Gene enrichment and depletion analysis of gemcitabine and selinexor screens.

**Table S1A.** Gene ranking according to β scores in gemcitabine and selinexor screens.

**Table S1B.** Lists of genes enriched in both screens.

**Table S1C.** Lists of genes depleted in both screens.

**Table S1D.** GO term enrichment analysis of enriched genes.

**Table S1E.** GO term enrichment analysis of depleted genes.

**Table S2.** Transcriptomic analysis of control, persisters, and recovered cells.

**Table S2A.** Differential gene expression between control cells and persisters.

**Table S2B.** Differential gene expression between control and recovered cells.

**Table S3.** mTOR-regulated persister signature in human patients’ residual tumors.

**Table S3A.** Genes included in the mTOR-regulated persister signature.

**Table S3B.** NES, norminal p, and FDR q values for GSEA of public data sets.

**Table S4.** Chemical library screen to identify mechanisms of persister survival.

**Table S4A.** List of small-molecule chemicals included in the library screen.

**Table S4B.** Chemical inhibitors significantly decreasing or increasing persisters.

## Notes

### Competing Interest Statement

The authors have declared no competing interest.

