## Supplementary Figure S1-S8 for "mTOR is a Major Determinant of Chemosensitivity"

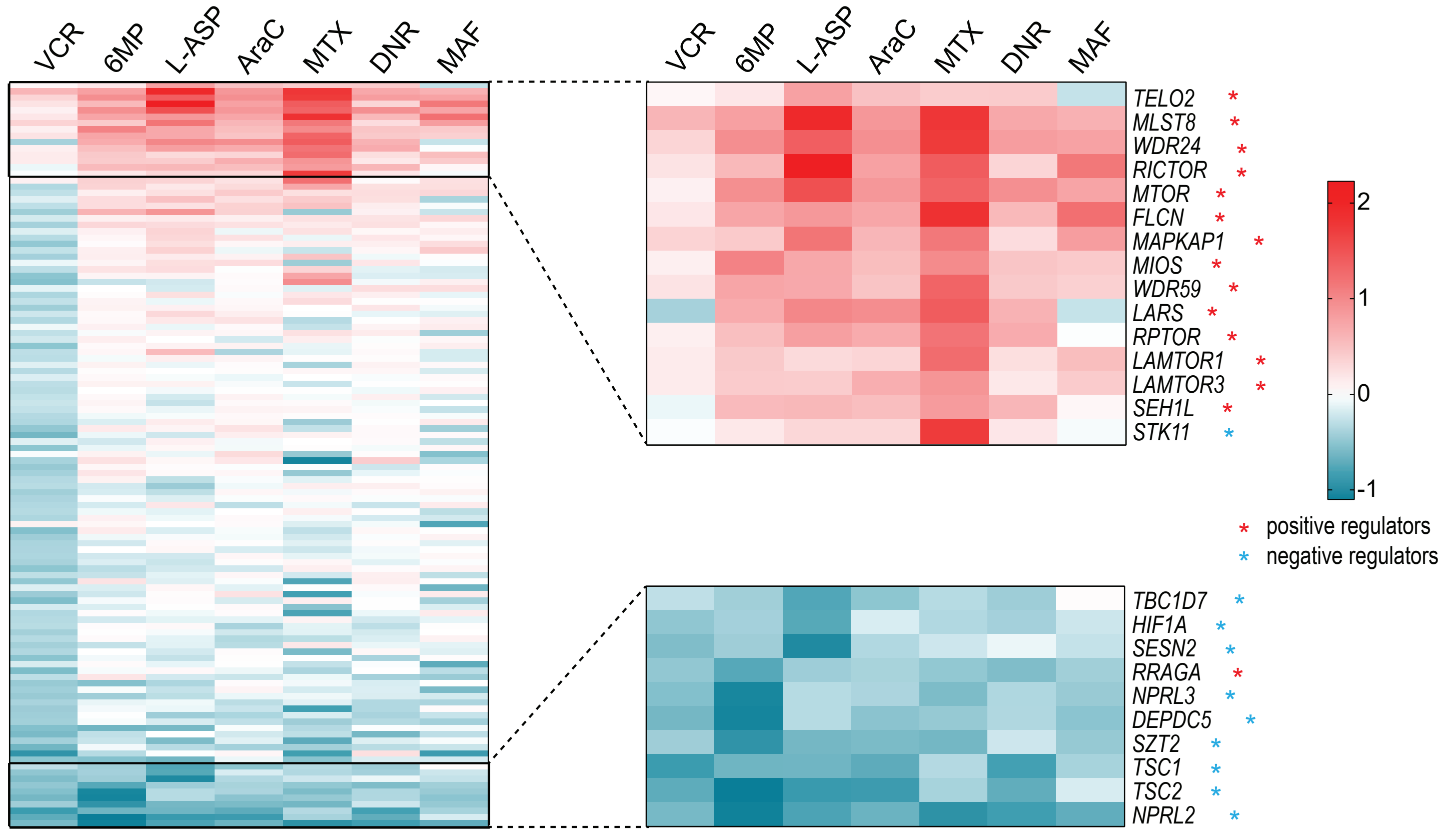
VCR
6MP
L-ASP
AraC
MTX
DNR
MAF
TELO2 *
MLST8 *
WDR24 *
RICTOR *
MTOR *
FLCN *
MAPKAP1 *
MIOS *
WDR59 *
LARS *
RPTOR *
LAMTOR1 *
LAMTOR3 *
SEH1L *
STK11 *
TBC1D7 *
HIF1A *
SESN2 *
RRAGA *
NPRL3 *
DEPDC5 *
SZT2 *
TSC1 *
TSC2 *
NPRL2 *
2
1
0
-1
* positive regulators
* negative regulators

### Figure S2

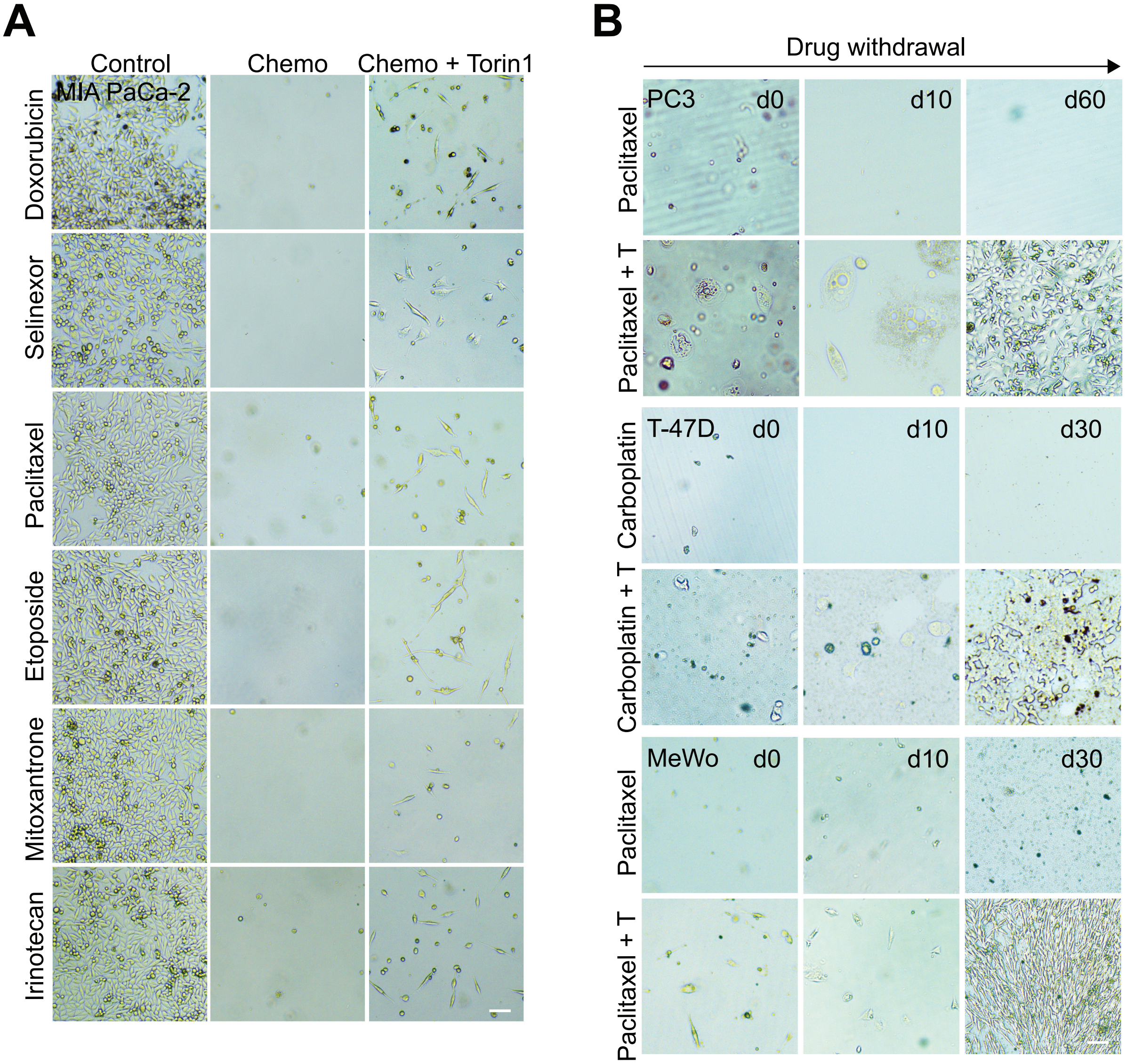

A
Control
Chemo
Chemo + Torin1
MIA PaCa-2
Doxorubicin
Selinexor
Paclitaxel
Etoposide
Mitoxantrone
Irinotecan
B
Drug withdrawal
PC3
d0
d10
d60
Paclitaxel
Paclitaxel + T
T-47D
d0
d10
d30
Carboplatin
Carboplatin + T
MeWo
d0
d10
d30
Paclitaxel
Paclitaxel + T

### Figure S2 *(continued)*

## C

| Pancreatic cancer | P53 status | Gemcitabine | Irinotecan | Etoposide |
|---|---|---|---|---|
| MDA-Panc-28 | Mutant | Yes | Yes | Yes |
| BxPC-3 | Mutant | Yes | Yes | Yes |
| MPanc96 | Mutant | No | Yes | Yes |
| Capan-2 | WT | Yes | No | No |
| **Prostate cancer** | **P53 status** | **Paclitaxel** | **Mitoxantrone** | **Estramustine** |
| LNCaP | Mutant | Yes | n.d | No |
| C4-2B | Mutant | Yes | Yes | Yes |
| DU145 | Mutant | Yes | Yes | Yes |
| 22Rv1 | Mutant | Yes | No | Yes |
| PC3 | Mutant | Yes | n.d | n.d |
| **Breast cancer** | **P53 status** | **Paclitaxel** | **Carboplatin** | **Gemcitabine** |
| T-47D | Mutant | Yes | Yes | Yes |
| MCF7 | WT | No | No | No |
| MDA-MB-231 | Mutant | Yes | Yes | Yes |
| SK-BR-3 | Mutant | Yes | Yes | Yes |
| **Liver cancer** | **P53 status** | **Gemcitabine** | **Doxorubicin** | **Mitoxantrone** |
| PLC/PRF/5 | Mutant | Yes | Yes | n.d |
| SNU398 | Mutant | Yes | Yes | Yes |
| Hep3B | Mutant | No | No | No |
| HepG2 | WT | No | No | No |
| **Melanoma** | **P53 status** | **Paclitaxel** | **Carboplatin** | **Gemcitabine** |
| Mewo | Mutant | Yes | Yes | Yes |
| A375 | WT | No | n.d | No |
| NM2C5 | Mutant | Yes | Yes | Yes |
| **Colon cancer** | **P53 status** | **Irinotecan** | **Gemcitabine** | **Etoposide** |
| HCT116 | WT | n.d | No | No |
| SW480 | Mutant | Yes | Yes | Yes |
| **Lung cancer** | **P53 status** | **Etoposide** | **Gemcitabine** | **Irinotecan** |
| H460 | WT | No | No | No |
| A549 | WT | No | No | No |
| H1299 | Mutant | Yes | Yes | Yes |
| H441 | Mutant | Yes | Yes | Yes |
| **HNSCC** | **P53 status** | **Paclitaxel** | **Carboplatin** | **Gemcitabine** |
| SCC47 | WT | No | No | No |
| **Ovarian cancer** | **P53 status** | **Etoposide** | **Gemcitabine** | **Irinotecan** |
| SK-OV-3 | Mutant | Yes | Yes | Yes |
| **Cervical cancer** | **P53 status** | **Etoposide** | **Gemcitabine** | **Irinotecan** |
| HeLa | Mutant | Yes | n.d | Yes |
| **Osteosarcoma** | **P53 status** | **Etoposide** | **Gemcitabine** | **Doxorubicin** |
| U2OS | WT | Yes | Yes | Yes |

**Notes:**
Yes: Persisters induced by treatment with chemotherapeutic agents plus Torin1
No: No persisters observed following combined treatment
n.d: Not determined

### Figure S3

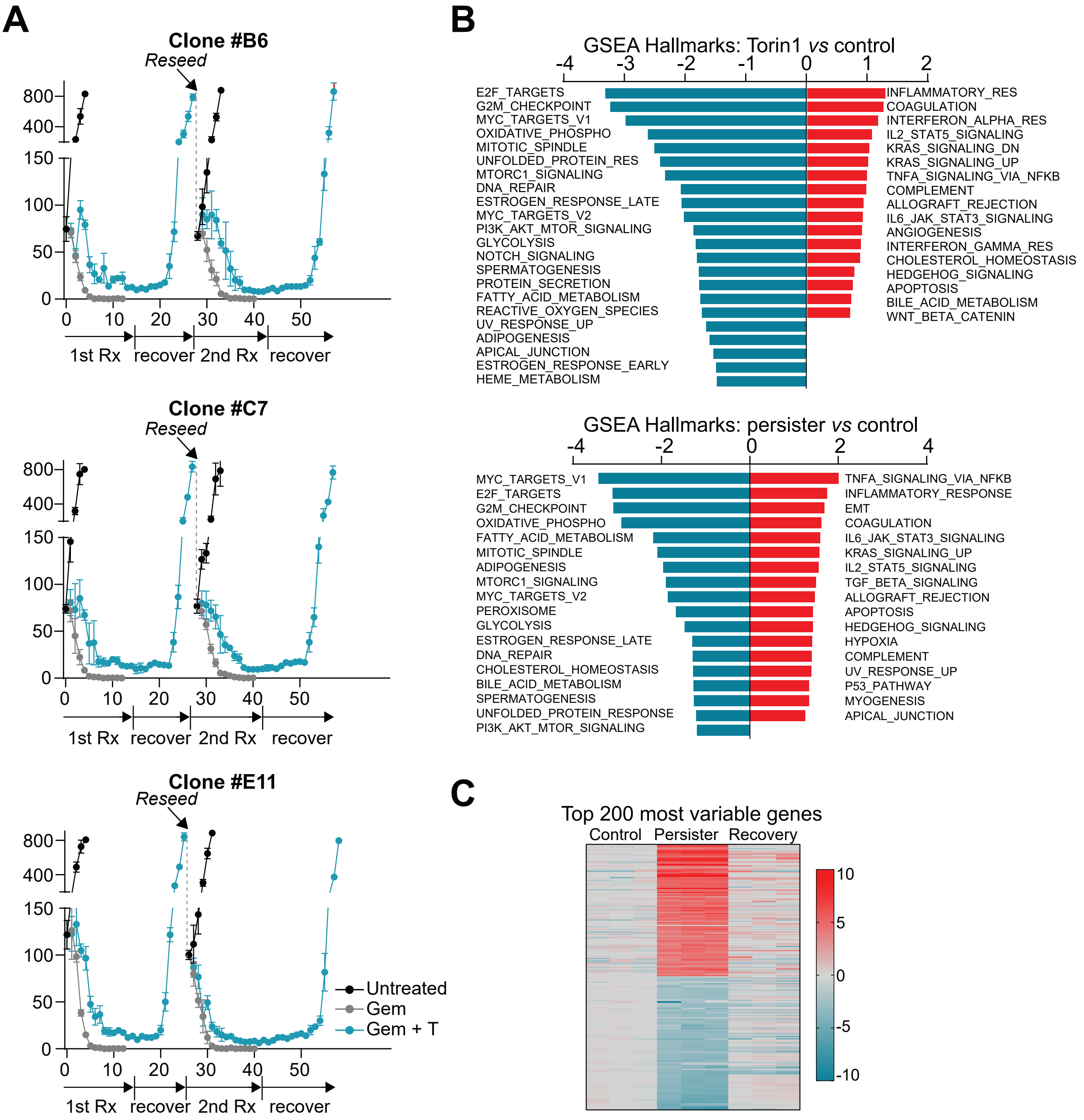
A
Clone #B6
Reseed
800
400
150
100
50
0
0 10 20 30 40 50
1st Rx
recover
2nd Rx
recover
Clone #C7
Reseed
Clone #E11
Reseed
Untreated
Gem
Gem + T
B
GSEA Hallmarks: Torin1 vs control
-4 -3 -2 -1 0 1 2
E2F_TARGETS
G2M_CHECKPOINT
MYC_TARGETS_V1
OXIDATIVE_PHOSPHO
MITOTIC_SPINDLE
UNFOLDED_PROTEIN_RES
MTORC1_SIGNALING
DNA_REPAIR
ESTROGEN_RESPONSE_LATE
MYC_TARGETS_V2
PI3K_AKT_MTOR_SIGNALING
GLYCOLYSIS
NOTCH_SIGNALING
SPERMATOGENESIS
PROTEIN_SECRETION
FATTY_ACID_METABOLISM
REACTIVE_OXYGEN_SPECIES
UV_RESPONSE_UP
ADIPOGENESIS
APICAL_JUNCTION
ESTROGEN_RESPONSE_EARLY
HEME_METABOLISM
INFLAMMATORY_RES
COAGULATION
INTERFERON_ALPHA_RES
IL2_STAT5_SIGNALING
KRAS_SIGNALING_DN
KRAS_SIGNALING_UP
TNFA_SIGNALING_VIA_NFKB
COMPLEMENT
ALLOGRAFT_REJECTION
IL6_JAK_STAT3_SIGNALING
ANGIOGENESIS
INTERFERON_GAMMA_RES
CHOLESTEROL_HOMEOSTASIS
HEDGEHOG_SIGNALING
APOPTOSIS
BILE_ACID_METABOLISM
WNT_BETA_CATENIN
GSEA Hallmarks: persister vs control
-4 -2 0 2 4
MYC_TARGETS_V1
E2F_TARGETS
G2M_CHECKPOINT
OXIDATIVE_PHOSPHO
FATTY_ACID_METABOLISM
MITOTIC_SPINDLE
ADIPOGENESIS
MTORC1_SIGNALING
MYC_TARGETS_V2
PEROXISOME
GLYCOLYSIS
ESTROGEN_RESPONSE_LATE
DNA_REPAIR
CHOLESTEROL_HOMEOSTASIS
BILE_ACID_METABOLISM
SPERMATOGENESIS
UNFOLDED_PROTEIN_RESPONSE
PI3K_AKT_MTOR_SIGNALING
TNFA_SIGNALING_VIA_NFKB
INFLAMMATORY_RESPONSE
EMT
COAGULATION
IL6_JAK_STAT3_SIGNALING
KRAS_SIGNALING_UP
IL2_STAT5_SIGNALING
TGF_BETA_SIGNALING
ALLOGRAFT_REJECTION
APOPTOSIS
HEDGEHOG_SIGNALING
HYPOXIA
COMPLEMENT
UV_RESPONSE_UP
P53_PATHWAY
MYOGENESIS
APICAL_JUNCTION
C
Top 200 most variable genes
Control
Persister
Recovery
10
5
0
-5
-10

### Figure S4

**A**

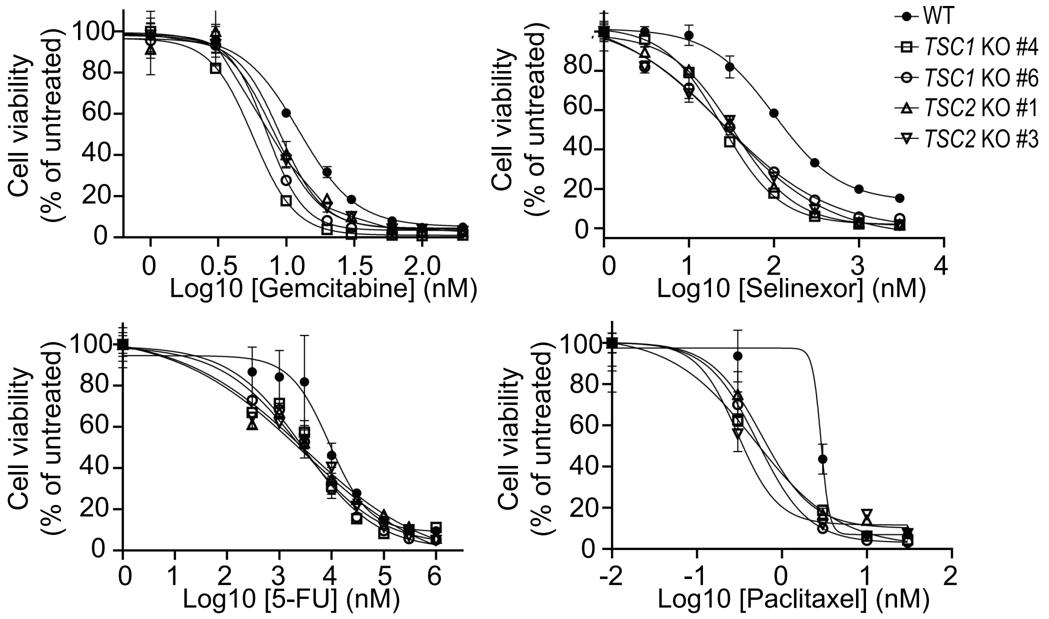

WT
TSC1 KO #4
TSC1 KO #6
TSC2 KO #1
TSC2 KO #3
Cell viability (% of untreated)
Log10 [Gemcitabine] (nM)
Log10 [Selinexor] (nM)
Log10 [5-FU] (nM)
Log10 [Paclitaxel] (nM)


**B**

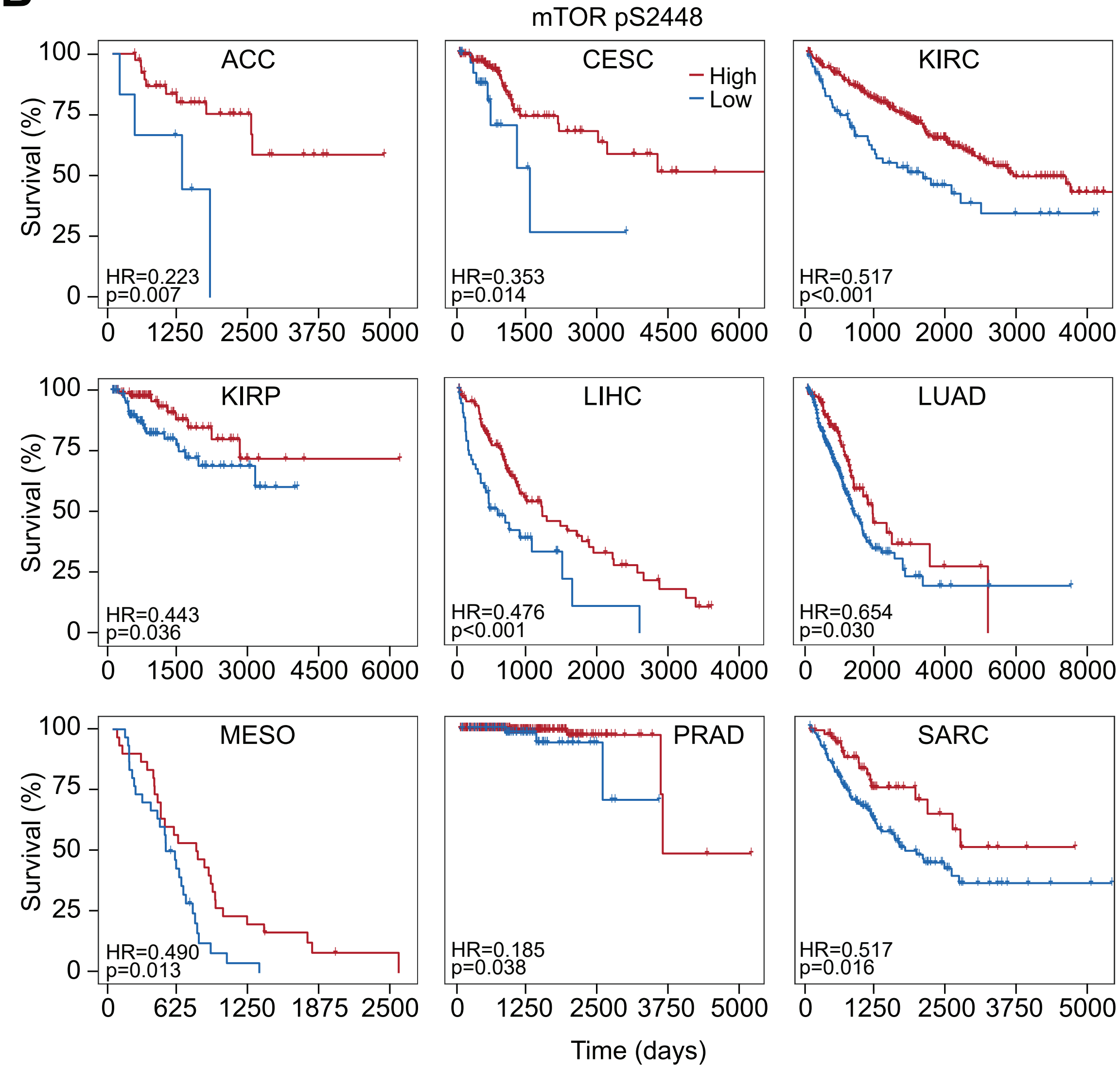

mTOR pS2448
ACC
HR=0.223
p=0.007
CESC
High
Low
HR=0.353
p=0.014
KIRC
HR=0.517
p<0.001
KIRP
HR=0.443
p=0.036
LIHC
HR=0.476
p<0.001
LUAD
HR=0.654
p=0.030
MESO
HR=0.490
p=0.013
PRAD
HR=0.185
p=0.038
SARC
HR=0.517
p=0.016
Survival (%)
Time (days)

### Figure S5

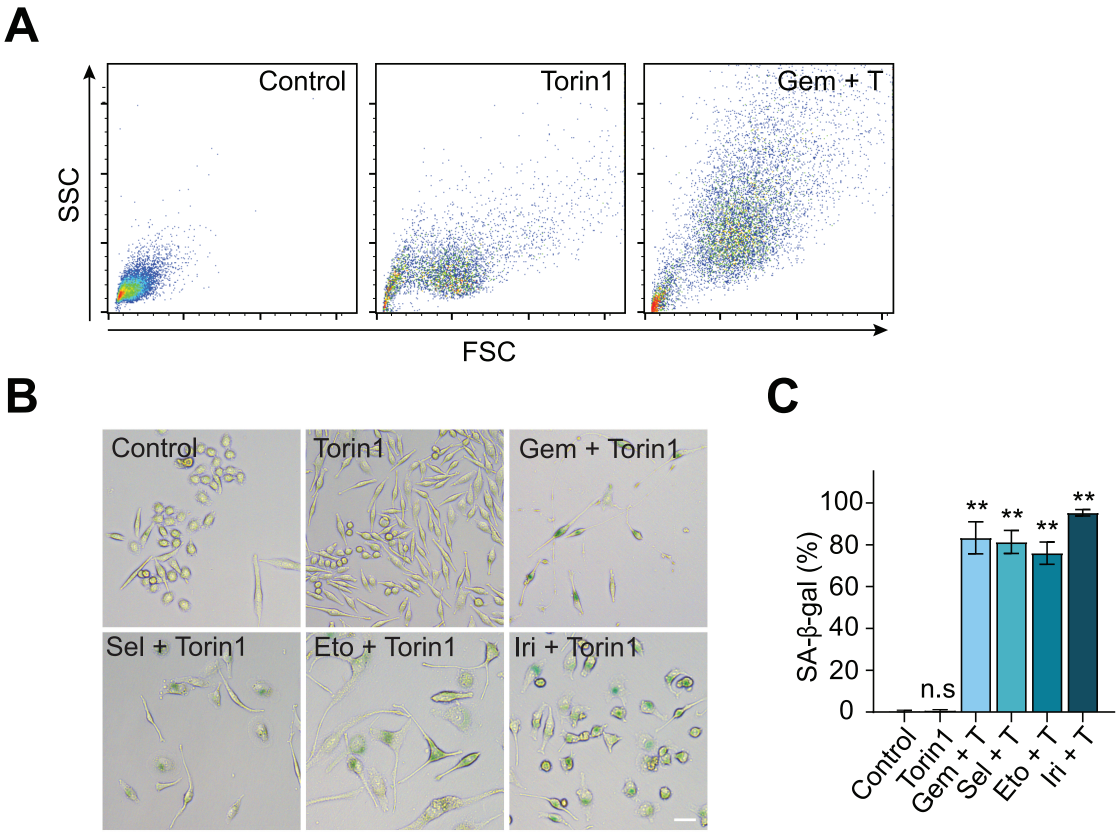
A
Control
Torin1
Gem + T
SSC
FSC
B
Control
Torin1
Gem + Torin1
Sel + Torin1
Eto + Torin1
Iri + Torin1
C
SA-β-gal (%)
100
80
60
40
20
0
n.s
**
Control
Torin1
Gem + T
Sel + T
Eto + T
Iri + T

### Figure S6

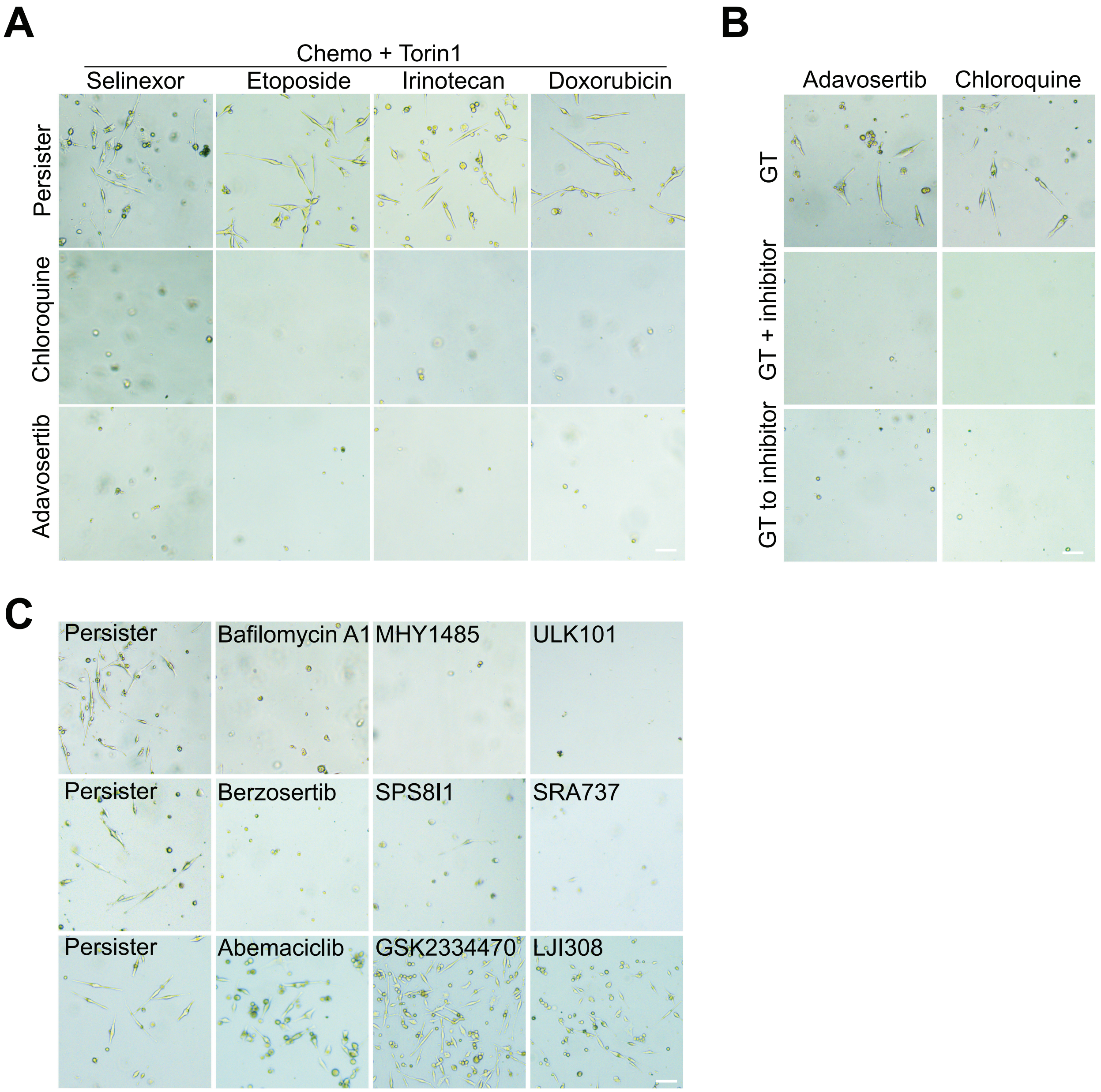
A
Chemo + Torin1
Selinexor
Etoposide
Irinotecan
Doxorubicin
Persister
Chloroquine
Adavosertib
B
Adavosertib
Chloroquine
GT
GT + inhibitor
GT to inhibitor
C
Persister
Bafilomycin A1
MHY1485
ULK101
Persister
Berzosertib
SPS8I1
SRA737
Persister
Abemaciclib
GSK2334470
LJI308

### Figure S7

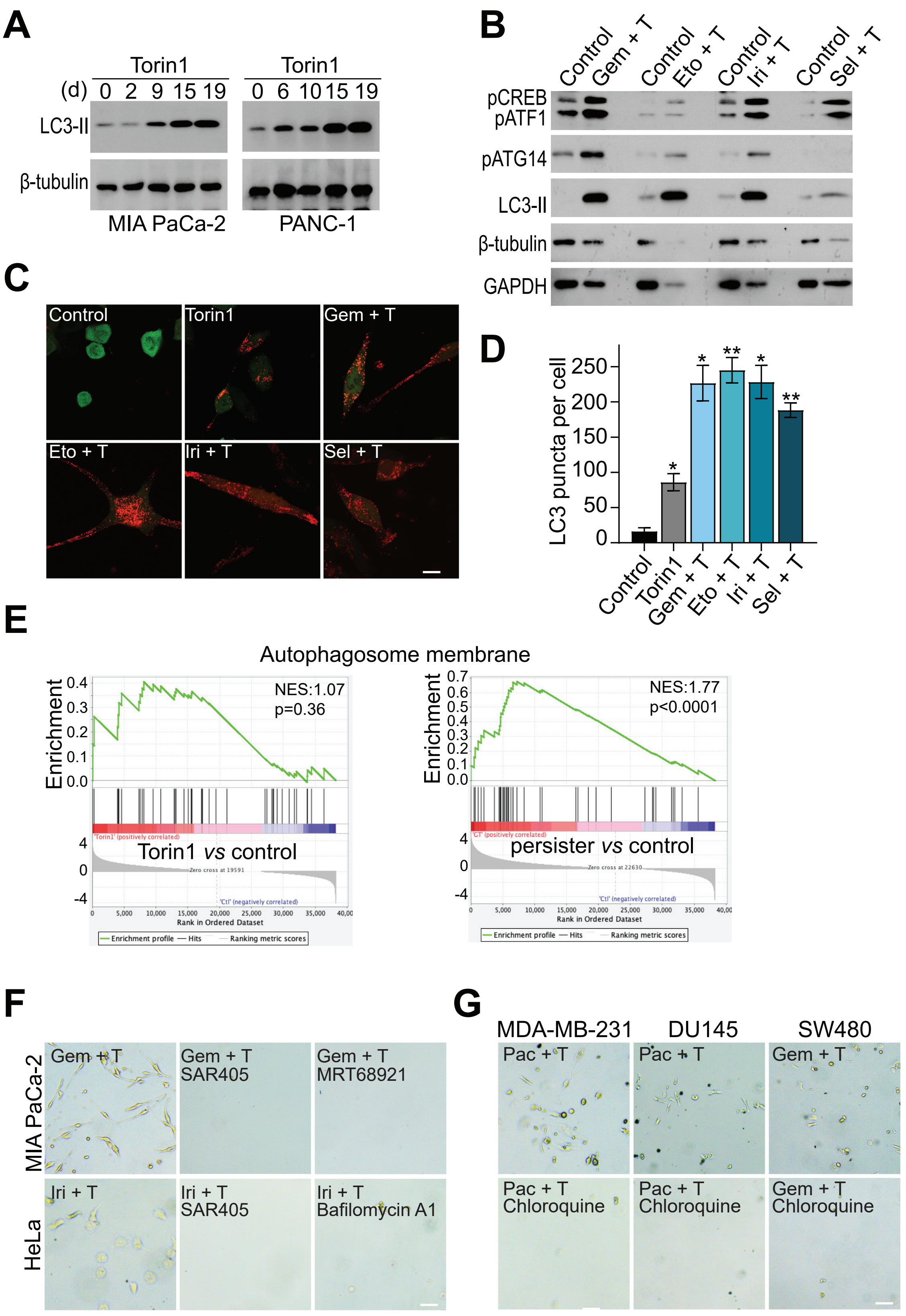

A
Torin1
Torin1
(d) 0 2 9 15 19
0 6 10 15 19
LC3-II
β-tubulin
MIA PaCa-2
PANC-1
B
Control
Gem + T
Control
Eto + T
Control
Iri + T
Control
Sel + T
pCREB
pATF1
pATG14
LC3-II
β-tubulin
GAPDH
C
Control
Torin1
Gem + T
Eto + T
Iri + T
Sel + T
D
LC3 puncta per cell
250
200
150
100
50
0
Control
Torin1
Gem + T
Eto + T
Iri + T
Sel + T
E
Autophagosome membrane
Enrichment
NES:1.07
p=0.36
Torin1 vs control
NES:1.77
p<0.0001
persister vs control
Rank in Ordered Dataset
F
MIA PaCa-2
HeLa
Gem + T
Gem + T SAR405
Gem + T MRT68921
Iri + T
Iri + T SAR405
Iri + T Bafilomycin A1
G
MDA-MB-231
DU145
SW480
Pac + T
Pac + T
Gem + T
Pac + T Chloroquine
Pac + T Chloroquine
Gem + T Chloroquine

### Figure S8

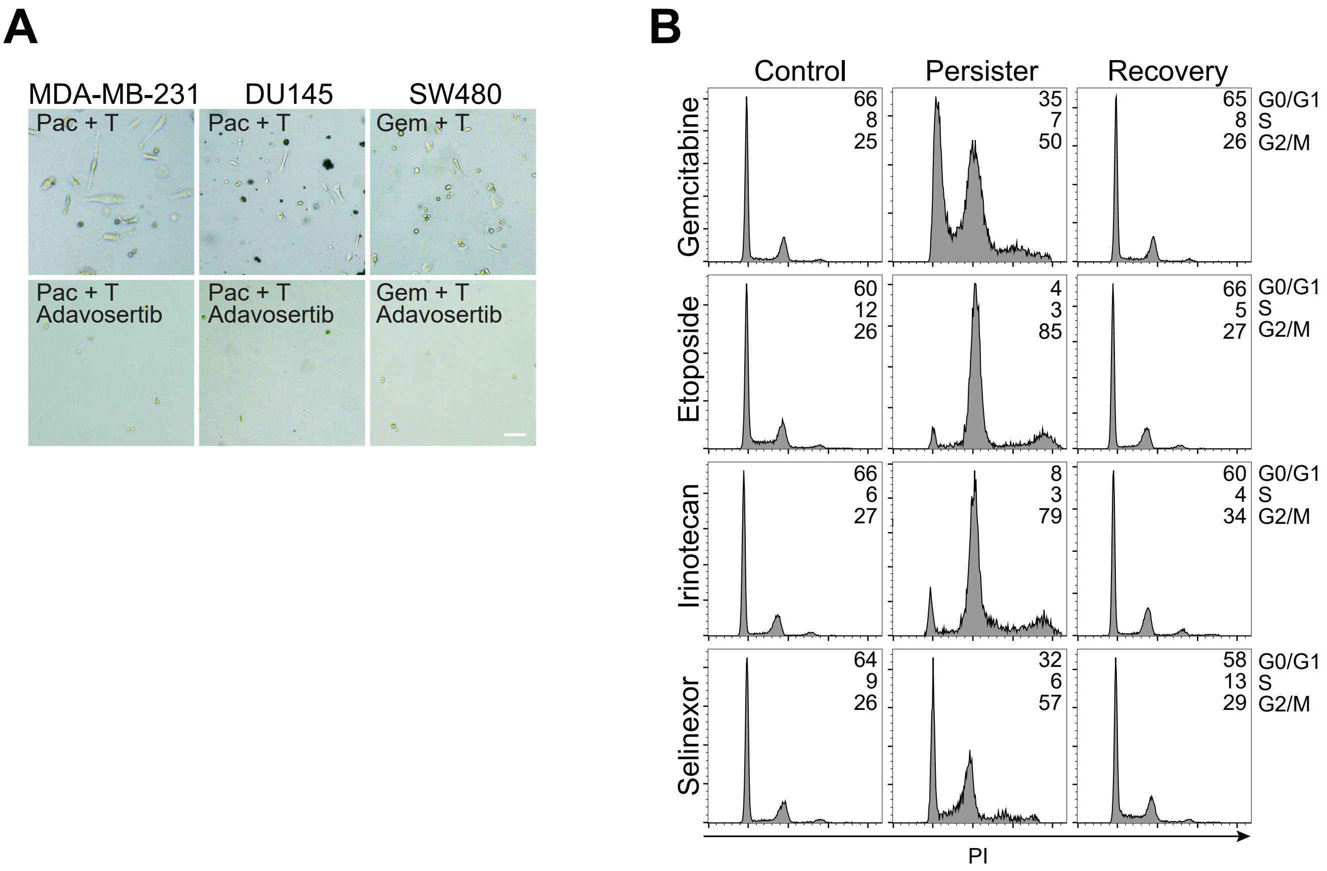
A
MDA-MB-231
DU145
SW480
Pac + T
Pac + T
Gem + T
Pac + T
Adavosertib
Pac + T
Adavosertib
Gem + T
Adavosertib
B
Control
Persister
Recovery
Gemcitabine
66
8
25
35
7
50
65
8
26
G0/G1
S
G2/M
Etoposide
60
12
26
4
3
85
66
5
27
G0/G1
S
G2/M
Irinotecan
66
6
27
8
3
79
60
4
34
G0/G1
S
G2/M
Selinexor
64
9
26
32
6
57
58
13
29
G0/G1
S
G2/M
PI
